# Diversified, miniaturized and ancestral parts for mammalian genome engineering and molecular recording

**DOI:** 10.1101/2024.09.30.615957

**Authors:** Troy A. McDiarmid, Megan L. Taylor, Wei Chen, Florence M. Chardon, Junhong Choi, Hanna Liao, Xiaoyi Li, Haedong Kim, Jean-Benoît Lalanne, Tony Li, Jenny F. Nathans, Beth K. Martin, Jordan Knuth, Alessandro L.V. Coradini, Jesse M. Gray, Sudarshan Pinglay, Jay Shendure

**Affiliations:** Department of Genome Sciences, University of Washington, Seattle, WA, USA; Seattle Hub for Synthetic Biology, Seattle, WA, USA; Developmental Biology Program, Memorial Sloan Kettering Cancer Center, New York, NY, USA; Brotman Baty Institute for Precision Medicine, Seattle, WA, USA; Howard Hughes Medical Institute, Seattle, WA, USA; Allen Discovery Center for Cell Lineage Tracing, Seattle, WA, USA

## Abstract

As the synthetic biology and genome engineering fields mature and converge, there is a clear need for a “parts list” of components that are diversified with respect to both functional activity (to facilitate design) and primary sequence (to facilitate assembly). Here we designed libraries composed of extant, ancestral, mutagenized or miniaturized variants of Pol III promoters or guide RNA (gRNA) scaffolds and quantified their ability to mediate precise edits to the mammalian genome via multiplex prime editing. We identified thousands of parts that reproducibly drive a range of editing activities in human and mouse stem cells and cancer cell lines, including hundreds exhibiting similar or greater activity than the sequences used in conventional genome engineering constructs. We further conducted saturation mutagenesis screens of canonical Pol III promoters (U6p, 7SKp, H1p) and the prime editing guide RNA (pegRNA) scaffold, which identified tolerated variants that can be superimposed on baseline parts to further enhance sequence diversity. While characterizing thousands of orthologous promoters from hundreds of extant or ancestral genomes, we incidentally mapped the functional landscape of mammalian Pol III promoter evolution. Finally, to showcase the usefulness of these parts, we designed a “ten key” molecular recording array that lacks repetitive subsequences in order to facilitate its one-step assembly in yeast. Upon delivering this 15.8 kb tandem array of promoters and guides to mammalian cells, individual pegRNAs exhibited balanced activities as predicted by the activity of component parts, despite their relocation to a single locus. Looking forward, we anticipate that the diversified parts and variant effect maps reported here can be leveraged for the design, assembly and deployment of synthetic loci encoding arrays of gRNAs exhibiting predictable, differentiated levels of activity, which will be useful for multiplex perturbation, advanced biological recorders and complex genetic circuits.

## Introduction

A central goal of synthetic biology is the design, synthesis and deployment of complex genetic circuits that measure and/or manipulate biological systems^1–8^. The components used in such circuits are often described as a “parts list”, wherein each “part” behaves and interacts with other exogenous parts (or endogenous factors) in a predictable manner, analogous to the parts lists of other engineering disciplines, *e.g.* the resistors, capacitors and inductors of electrical circuits^9,10^. However, a unique challenge for synthetic biology is that its parts, typically encoded as DNA, are unstable if repetitive, *i.e.* if the same subsequence appears repeatedly in different parts that are used in the same *cis-*encoded circuit. The challenges associated with repetitive subsequences manifest at nearly every step, but are most problematic during synthesis and assembly^11–14^. For example, although yeast-based assembly can now be used to construct entirely synthetic loci that are over 100 kilobases^15–18^ in length, the homologous recombination (HR) mechanisms that enable yeast-based assembly also corrupt the process if the same subsequence appears repeatedly. As a consequence, the same part cannot easily be used more than once in a yeast-assembled, single-locus, mammalian-deployed genetic circuit.

This practical challenge is particularly evident at the intersection of synthetic biology and genome engineering. For example, we and others have envisioned multiplex cell lineage recorders that rely on many instances of Pol III promoters, gRNAs and target sites, ideally encoded at a single locus to facilitate the generation of distributable “recorder cell lines” and “recorder mice”^19–23^. However, the number of sequence-diverse “parts” that are validated and characterized for these functionalities is very limited, precluding the assembly of synthetic loci with tens to hundreds of Pol III promoter-gRNA units. For example, for Pol III transcription, the field overwhelmingly relies on a handful of endogenous human promoters (usually U6, sometimes H1 or 7SK), and for gRNA scaffolds, on one of two designs derived from *S. pyogenes*^5,24–28^. Although substantial progress has been made towards the design and identification of non-repetitive parts for CRISPR genome engineering that are operational in <u>bacterial</u> systems^29,30^, we continue to lack validated, diversified parts with well-characterized functionality for genome engineering of <u>mammalian</u> systems.

In this study, we sought to address this by first designing diverse libraries of Pol III promoters and gRNA scaffolds, and then quantifying their activity with a multiplex prime editing-based functional assay. Through these experiments, we validate and characterize thousands of sequence-diverse parts capable of driving genome editing in human and mouse cancer and stem cell lines. Both Pol III promoter and gRNA scaffold variants exhibited highly reproducible activities spanning several orders of magnitude, including parts that are more compact and/or more active than the most widely used sequences. Finally, we demonstrate how these diversified promoters and gRNA scaffolds can be leveraged to design multi-component synthetic loci that are easily assembled in yeast. Specifically, we design and assemble a single locus, 10-key diversified molecular recording array^31^, and demonstrate that its tandemly arranged parts function as predicted in mammalian cells.

## Results

### Design, synthesis and functional characterization of diversified U6 promoters

To date, only a handful of Pol III promoters have been characterized for genome engineering in mammalian cells^24,28,32^. To remedy this, we designed a library of 209 diversified Pol III U6 promoters through several complementary approaches (**Fig. 1a**; **Table S1**). A first subset are “evolutionarily diversified” U6 promoters (n = 97, promoter length range: 249-600 bp, mean length = 475 bp). These include 89 orthologs of human U6 promoters with putative transcriptional activity^33^ from across vertebrate species, the canonical human RNU6-1 promoter that is widely used in mammalian RNAi and gRNA delivery vectors^24,34–37^, 4 mammalian promoters designed for a 3-gRNA array lentiviral Perturb-seq vector^24,34–37^, and finally 3 additional human U6 promoters that were sufficiently divergent from the human RNU6-1 promoter^33^. A second subset are “synthetically diversified” U6 promoters (n = 112, promoter length range: 249-252 bp, mean length = 250 bp). We designed these by shuffling the nucleotides located in between known core transcription factor binding sites (TFBS) while also introducing putatively tolerated SNVs to TFBS; the template for this programmed diversification was the human RNU6-1 promoter.

**Figure 1.**
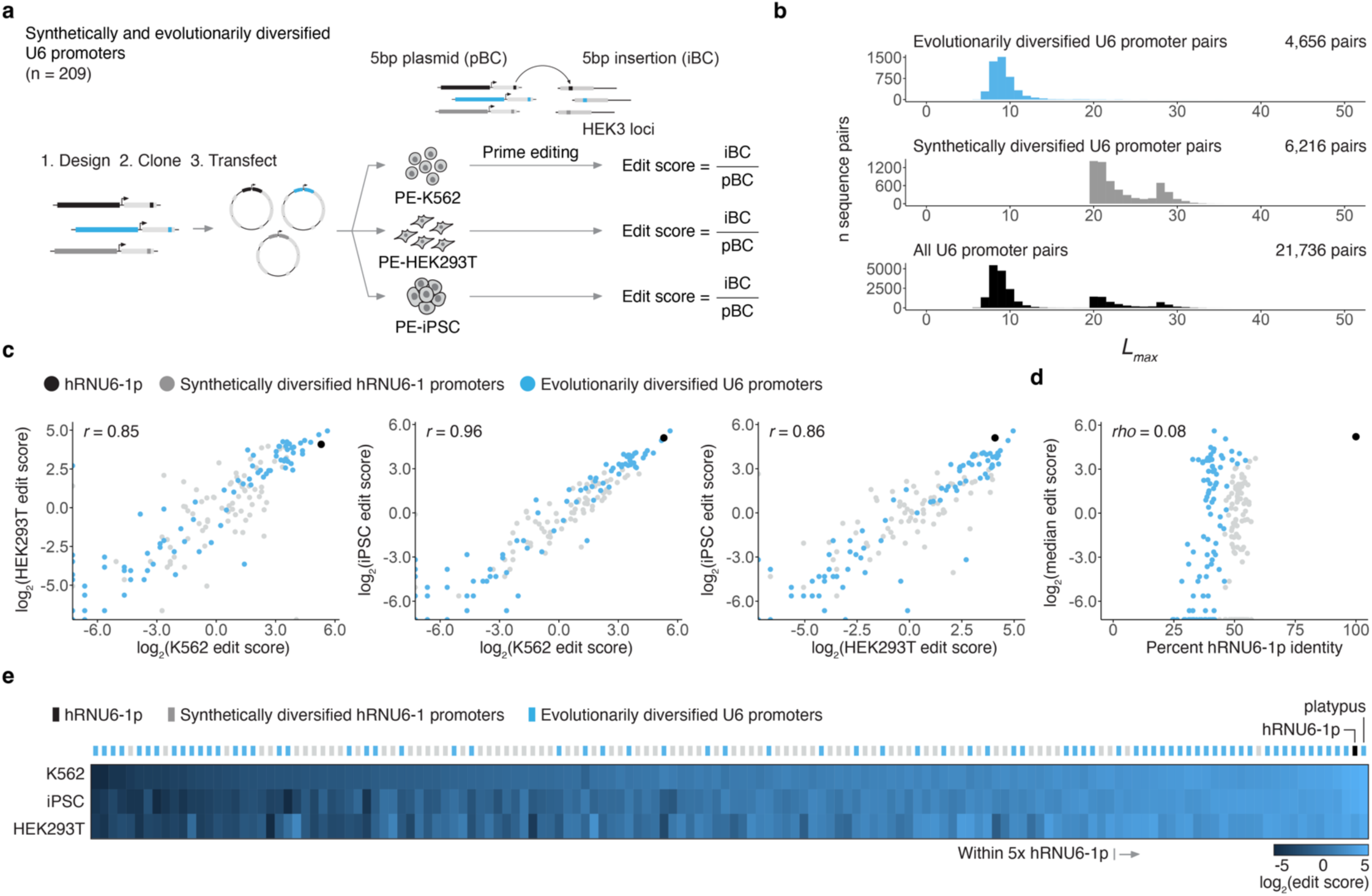
Multiplex functional characterization of synthetically and evolutionarily diversified U6 promoters in human cells. **a)** Synthetically and evolutionarily diversified U6 promoters were tested in three human cellular contexts with a multiplex prime editing functional assay. Edit scores were defined as the frequency of an insertional barcode (iBC) at the genomic target site divided by the frequency of the same barcode in the plasmid library (pBC). **b)** *L_max_* distributions quantifying the maximal shared repeat length between all possible pairs of sequences for the evolutionarily diversified U6 promoter library (n = 97; 4,656 pairs), the synthetically diversified hRNU6-1p library (n = 112; 6,216 pairs) and the combined set (n = 209; 21,736 pairs) in the same orientation. See **Supplementary Fig. 1b** for *L_max_* distributions for reverse complement comparisons. **c)** Pairwise comparison of log-transformed edit scores between cellular contexts. Pearson correlations, calculated on barcode-normalized edit scores prior to log transformation, are shown. **d)** Sequence identity with hRNU6-1p (*x-*axis) is not predictive of functional activity of synthetically or evolutionarily diversified U6 promoters. Spearman correlation is shown. **e)** Edit scores of 146 functional diversified U6 promoters ordered left-to-right by ascending median edit score across three human cellular contexts.

To quantify and ensure sequence diversity, we developed an algorithm that calculates the length and identity of the longest shared repeat between every possible pair of sequences in either orientation, termed *L_max_* (**Supplementary Fig. 1**)^29^. Applying this algorithm to the set of 209 diversified Pol III U6 promoters, we found that they can be used in any of the 21,736 possible pairwise combinations and satisfy *L_max_* < 40, a practical requirement for ensuring compatibility with contemporary protocols for large-scale assembly of synthetic DNA in yeast (*L_max_* for the full set = 36; **Fig. 1b**; **Supplementary Fig. 1**)^15,17^.

We then sought to perform a multiplex experiment that quantified the relative activity of these Pol III promoters. For this, we cloned the promoters upstream of a pegRNA designed to install a 5 bp insertional barcode at the *HEK3* locus in the human genome, with a strategy that linked each Pol III promoter to a specific barcode (**Fig. 1a**)^38,39^. In the experiments described below, we quantify the functional activity of a given promoter as the frequency of its insertional barcode at the genomic target site (iBC) normalized by the frequency of the same barcode in the plasmid library encoding the promoter-pegRNA combinations (pBC). We refer to this ratio as the edit score, analogous to regulatory element activity scores of massively parallel reporter assays (**Fig. 1a**)^39,40^. As the barcodes themselves influence editing efficiency^31^, we also measured the insertion efficiency of every possible 5N insertion (n = 1,024 barcodes) when driven by the standard human RNU6-1 promoter, and used this data to further normalize edit scores (**Supplementary Fig. 2**).

We introduced this library of Pol III promoter-driven pegRNAs to human K562 cells, HEK293T cells, or iPSCs that had been engineered to stably express a prime editor^38,41^. Both synthetically and evolutionarily diversified U6 promoters drove genome editing at the *HEK3* locus at a broad range of levels (**Fig. 1c-e**; **Supplementary Fig. 3**; **Table S1**). Edit scores were reasonably well correlated between technical replicates (*r* = 0.47-0.96) and cellular contexts *(r* = 0.85-0.96; **Fig. 1c**; **Supplementary Fig. 3**). Of note, evolutionarily diversified U6 promoters displayed greater variance in activity levels than synthetically diversified alternatives, consistent with their greater sequence divergence from the human RNU6-1 promoter (**Fig. 1c-e**; **Supplementary Fig. 4**). The canonical human RNU6-1 promoter was consistently among the most active promoters, modestly outperformed by only a U6 promoter of *Ornithorhynchus anatinus*, the duck-billed platypus (1.2-1.8-fold; **Fig. 1e**; **Table S1**).

Altogether, we identified 146/209 (70%) promoters that drove editing in all 3 cellular contexts (**Fig. 1e**). There were 70 promoters displaying edit scores > 1 across all contexts, which corresponds to activity within about 50-fold of the standard human RNU6-1 promoter (**Table S1**). Among these, there were 28 whose activity fell within 5-fold of the standard human RNU6-1p in all three contexts, including all three other human U6 promoters tested^33^, 2/4 promoters previously tested by Adamson et al.^24^, and 23 newly characterized promoters (21 evolutionary diversified, 2 synthetically diversified) (**Fig. 1e**; **Table S1**). Four of these 23 highly functional, newly characterized U6 promoters ranked higher than previously characterized non-human RNU6-1p orthologs, specifically those of the common snapping turtle (*Chelydra serpentina*), the one-humped camel (*Camelus dromedarius*), the domestic muscovy duck (*Cairina moschata domestica*), and finally the aforementioned platypus (**Fig. 1e**; **Table S1**).

We sought to validate these results using two strategies. First, we identified a subset of the diversified U6 promoters representing a broad range of activity levels in the primary screen and then re-cloned and independently tested them in a monoclonal PEmax-iPSC line (n = 50 diversified U6 promoters together with the standard human RNU6-1p). Results for this validation set correlated strongly with results from the primary screen (*r* = 0.93; **Supplementary Fig. 5a**). Next, we re-cloned our 10 top-performing promoters (standard human RNU6-1p and 9 evolutionarily diversified promoters) with alternate iBCs and again tested their activity in PEmax-iPSCs (5 alternate iBCs for human and platypus and 1 alternate iBC for the remaining eight promoters). All 10 of these top-performing U6 promoters drove robust editing to within 2.6-fold of the human RNU6-1p standard (**Supplementary Fig. 5b-c**). In these experiments, the platypus U6 promoter ortholog was once again a top-ranked promoter, albeit on par with rather than exceeding the activity of human RNU6-1p (**Supplementary Fig. 5b-c**).

Together with the primary screen, these results confirm synthetically and evolutionarily diversified U6 promoters from across species are functional in human cells and reproducibly exhibit a broad range of activities in driving genome editing. Although both strategies yielded functional promoters with activities within 5-fold of that of human RNU61-p, the vast majority of this highly active subset were evolutionarily diversified. Interestingly, while human RNU6-1p was consistently among the top performers in human cells, there were a few U6 promoters from extant species that exhibited comparable activity in human cells, despite extensive sequence divergence.

### Design, synthesis and functional characterization of diversified pegRNA scaffolds

Diversifying gRNA scaffolds is considerably more challenging than diversifying Pol III promoters due to extensive constraints on gRNA secondary structure^25,29,42–44^. We designed libraries of diversified pegRNA scaffolds to satisfy *L_max_* < 40 using two approaches. First, we introduced putatively secondary structure-retaining 5N and 4N replacements to repeat:anti-repeat (R:AR) regions (“replacement designs”). Second, we introduced 5N insertions to regions predicted to tolerate insertions based on pegRNA secondary structure, along with R:AR 5N replacements (“extension designs”). Altogether, we designed 174 replacement and 138 extension pegRNA scaffolds, and then specific versions of these to install a 5 bp insertional barcode (iBC) at the human *HEK3* locus (**Fig. 2a**; **Supplementary Fig. 6**).

**Figure 2.**
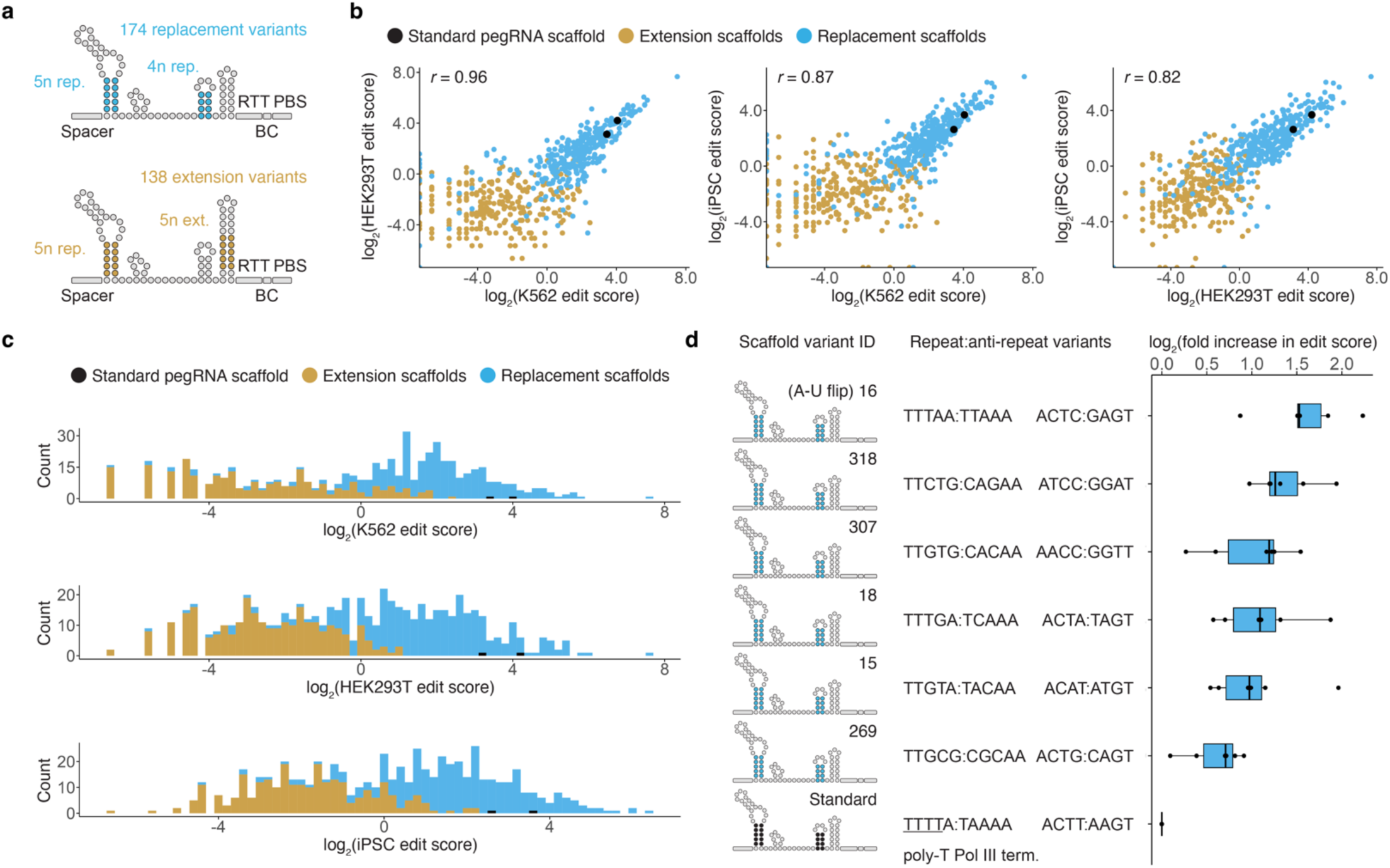
Multiplex functional characterization of diversified pegRNA scaffolds in human cells. **a)** Diversified pegRNA scaffold designs. Complementary R:AR sequences were introduced at specific locations, producing either replacement (top) or extension (bottom) variants of the conventional pegRNA scaffold. **b)** Pairwise comparison of log-transformed edit scores between cellular contexts. Pearson correlations, calculated on barcode-normalized edit scores prior to log transformation, are shown. **c)** Replacement scaffolds tended to have higher edit scores than extension scaffolds. **d)** Diversified pegRNA scaffolds that eliminated a Pol III termination sequence consistently exhibited higher edit scores than the standard scaffold. Boxes represent the 25th, 50th, and 75th percentiles. Whiskers extend from hinge to 1.5 times the interquartile range.

We synthesized and cloned these 312 pegRNA scaffold variants downstream of human RNU6-1p, each driving a specific iBC, and introduced them to human K562 cells, HEK293T cells, or iPSCs that stably expressed a prime editor^38,41^. Because the impact of the iBC sequence on pegRNA secondary structure and insertion efficiency can be difficult to predict^39,45^, we also synthesized and cloned each pegRNA scaffold with an alternate iBC in a second library, which was tested independently. After sequencing 5 bp insertional barcodes at the *HEK3* locus, we quantified the edit score for each scaffold-iBC pair and normalized these for differential iBC efficiencies as above (**Table S2**). Results correlated reasonably well across cellular contexts (*r* = 0.82-0.96; **Fig. 2b**; **Supplementary Fig. 7**) as well as across independent iBC sets (*r* = 0.53-0.85; **Supplementary Fig. 8**). Overall, replacement designs markedly outperformed insertion designs (13.2- to 36.0-fold higher median edit score across cellular contexts; **Fig. 2c**).

Altogether, we identified 267/312 (86%) pegRNA scaffolds that drove editing with both iBCs across all cellular contexts (**Table S2**). Among these, 56 functioned within 5-fold of the standard pegRNA scaffold with both iBCs across all cellular contexts, including six that outperformed the standard pegRNA scaffold (**Fig. 2d**; **Table S2**). These six included a scaffold with a previously described A-U flip design that swaps nucleotides in the first R:AR region to remove a polythymidine Pol III termination sequence (“TTTTA:TAAAA”>”TTTAA:TTAAA”), previously reported to improve function by reducing premature termination of Pol III transcription^26,46^. The remaining five scaffolds that outperformed the standard pegRNA each maintain the first two “TT” nucleotides in the first R:AR sequence while introducing variants that disrupt the Pol III termination sequence through means other than the A-U flip (**Fig. 2d**). Taken together, these results identify dozens of sequence-diversified pegRNA scaffolds that are similarly active to the conventional scaffold in human cells, and confirm two strategies to diversify (pe)gRNA scaffolds while maintaining or improving their function: namely introducing complementary R:AR variants and/or removing Pol III termination sequences.

### Saturation mutagenesis and functional assessment of a miniaturized U6p-pegRNA cassette

The diversified parts described thus far were designed to satisfy *L_max_* < 40, a practical requirement for yeast-based assembly of large constructs^15,17^. Smaller subsets of parts can be selected from these libraries to further increase diversity. However, more comprehensive knowledge regarding which variants can be introduced to a Pol III promoter and/or gRNA scaffold while retaining functionality would enable design of further diversified parts to satisfy even more stringent *L_max_* requirements. To this end, we conducted saturation mutagenesis and functional assessment of a U6p-pegRNA cassette.

To focus our efforts on the most critical sequence elements, our “wild-type” construct appends a miniaturized version of the canonical human RNU6-1 promoter^47^ that retains its four key TFBS while deleting divergent intervening regions (shortened from 249 to 111 bp; **Fig. 3a**; **Table S3**) to a standard pegRNA driving a 5 bp insertion (124 bp). We first sought to confirm that the wild-type version of this 235 bp minU6p-pegRNA cassette is functional, and found it drove editing at 38% of standard hRNU6-1p levels (**Fig. 3a**). In contrast, deletion of TFBS from minU6p severely diminished activity (169- to 2732-fold reduction; **Fig. 3a**). The H1 promoter, a naturally occurring human Pol III promoter, similarly miniaturized in the sense that the TFBS are retained, exhibited similar activity as miniaturized U6p (29% of standard hRNU6-1p; **Fig. 3a**). Taken together, these results confirm that retention of TFBS while deleting divergent intervening sequences is a general approach for deriving miniaturized Pol III promoters that retain function^47,48^.

**Figure 3.**
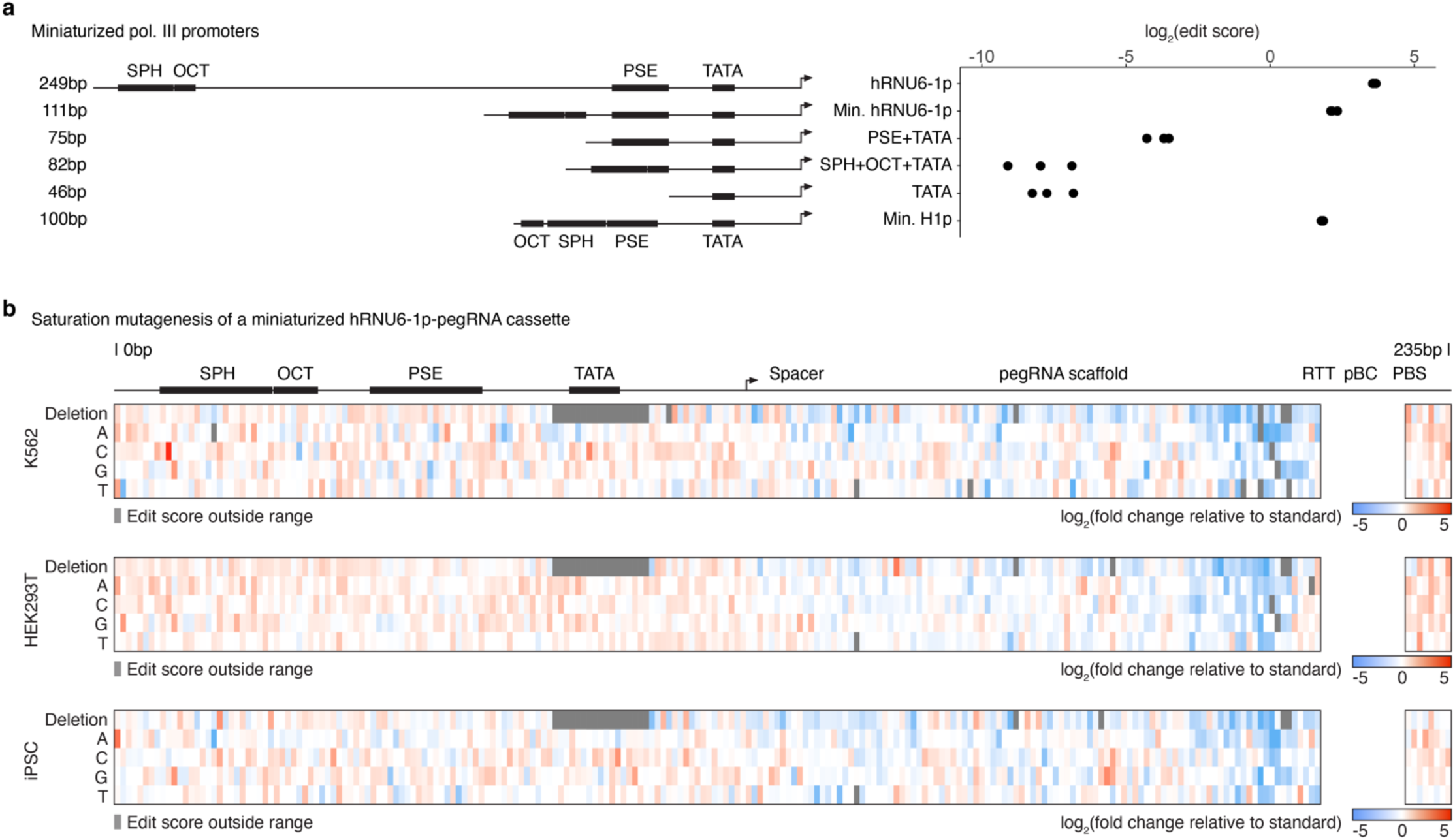
Saturation mutagenesis of a miniaturized U6p-pegRNA cassette. **a)** Left: Human Pol III promoter deletion series constructs and corresponding lengths. Locations of key TFBS are labeled. The top five rows correspond to hRNU6-1p and miniaturized variants thereof. The key TFBS are always in the same order from 5’ to 3’ (5’-SPH-OCT-PSE-TATA). The bottom row corresponds to the 100 bp human H1 promoter, in which the positions of the OCT and SPH elements are reversed relative to hRNU6-1p. Right: Log-scaled edit scores of wildtype or miniaturized Pol III promoters (n = 3 transfection replicates each with 4 iBCs per promoter, mean of the edit scores of these 4 iBCs per transfection replicate are shown). **b)** Variant effect maps of saturation mutagenesis of a miniaturized hRNU6-1p-pegRNA cassette tested across three human cellular contexts. Color-scaled, log-transformed fold-changes in median edit scores relative to minU6p-pegRNA are shown. Edit scores were not calculated for the unboxed region surrounding the pBC, as exact matches spanning this region were required for edit quantification.

With the wild-type miniaturized U6p-pegRNA as the baseline, we designed, synthesized and cloned two libraries encoding every possible single nucleotide substitution and and single nucleotide deletion across its length (230 bp excluding the 5N iBC; n = 920 variants in total; a second library is identical but with a different set of iBC pairings; **Table S3**). We then, as above, introduced these libraries to three human cellular contexts and quantified edit scores. These experiments revealed a biologically coherent landscape of variant effects with consistent sequence-function relationships across cellular contexts (**Fig. 3b**; **Supplementary Figs. S9**-**S10**). As expected given the flexibility of the *cis*-regulatory code, the U6 promoter region (positions 1-111) was more tolerant to variation than the pegRNA (positions 112-235; 1.6- to 1.9-fold higher median edit score across cell contexts; **Fig. 3b**; **Supplementary Fig. 10**). Single nucleotide deletions within the U6 promoter TATA box (positions 81-89: “TTTATATAT”) were not tolerated (**Fig. 3b**). Activity was also particularly compromised by deletions in the nucleotides forming the final pegRNA stem loop (positions 198-202: “GAGTC”; 2.1- to 5.4-fold lower edit scores than all other deletions) or PAM-proximal portion region of the spacer (positions 122-131: “GAGCACGTGA”; 1.4- to 1.6-fold lower edit scores than all other deletions; **Fig. 3b**; **Supplementary Fig. 10**). These results are consistent with the core roles of these elements in the editing cycle of a pegRNA: transcription, stability, and target nicking, respectively.

In contrast to single nucleotide deletions, many SNVs were tolerated throughout the length of the cassette, and several displayed enhanced performance compared to the miniaturized U6p-pegRNA cassette (**Fig. 3b**; **Table S3**). In particular, 16 of 920 variants, 15 of which were SNVs, displayed increased edit scores across both iBCs in all three cellular contexts (median 1.9-fold higher edit scores, max 20.8-fold; **Fig. 3b**; **Table S3**). 13/16 (81%) of these variants were in the miniaturized promoter, of which 5 introduced substitutions to a “TATT” sequence at the end of the proximal sequence element (PSE; positions 64-67), which may boost function by improving promoter conformation and/or transcription initiation from the immediately downstream TATA box. The 3/16 (19%) variants with improved function in the pegRNA region all introduced substitutions to two neighboring nucleotides near the 3’ end of the primer binding site (231G>C; 232T>C; 232T>A), suggesting these variants may yield a more optimal primer and/or more stable pegRNA. Relaxing these criteria, we identified 34 variants that functioned within 10% of the wild-type minU6p-pegRNA cassette across barcodes and contexts, and 214 that functioned within 2-fold. These results provide a rich set of enhancing or tolerated single nucleotide variants that can be leveraged to boost sequence diversity as needed (**Fig. 3b**; **Table S3**).

### Diversified U6 promoters exhibit consistent functional activities in mouse embryonic stem cells

To assess whether the activities of these parts are human-specific or consistent across mammalian models, we next sought to characterize them in mouse embryonic stem cells (mESCs). As mESCs lack an endogenous *HEK3* locus, we introduced synthetic human *HEK3* target sites^49^ (*synHEK3*) and PEmax via piggyBac transposition at a high multiplicity of integration, and isolated a monoclonal line with an estimated 87 *synHEK3* targets (29 integrations x 3 *synHEK3* targets per integration; **Fig. 4a**). We then introduced the original library of evolutionarily or synthetically diversified U6 promoters (n = 209) to this cell line and quantified edit scores as above.

**Figure 4.**
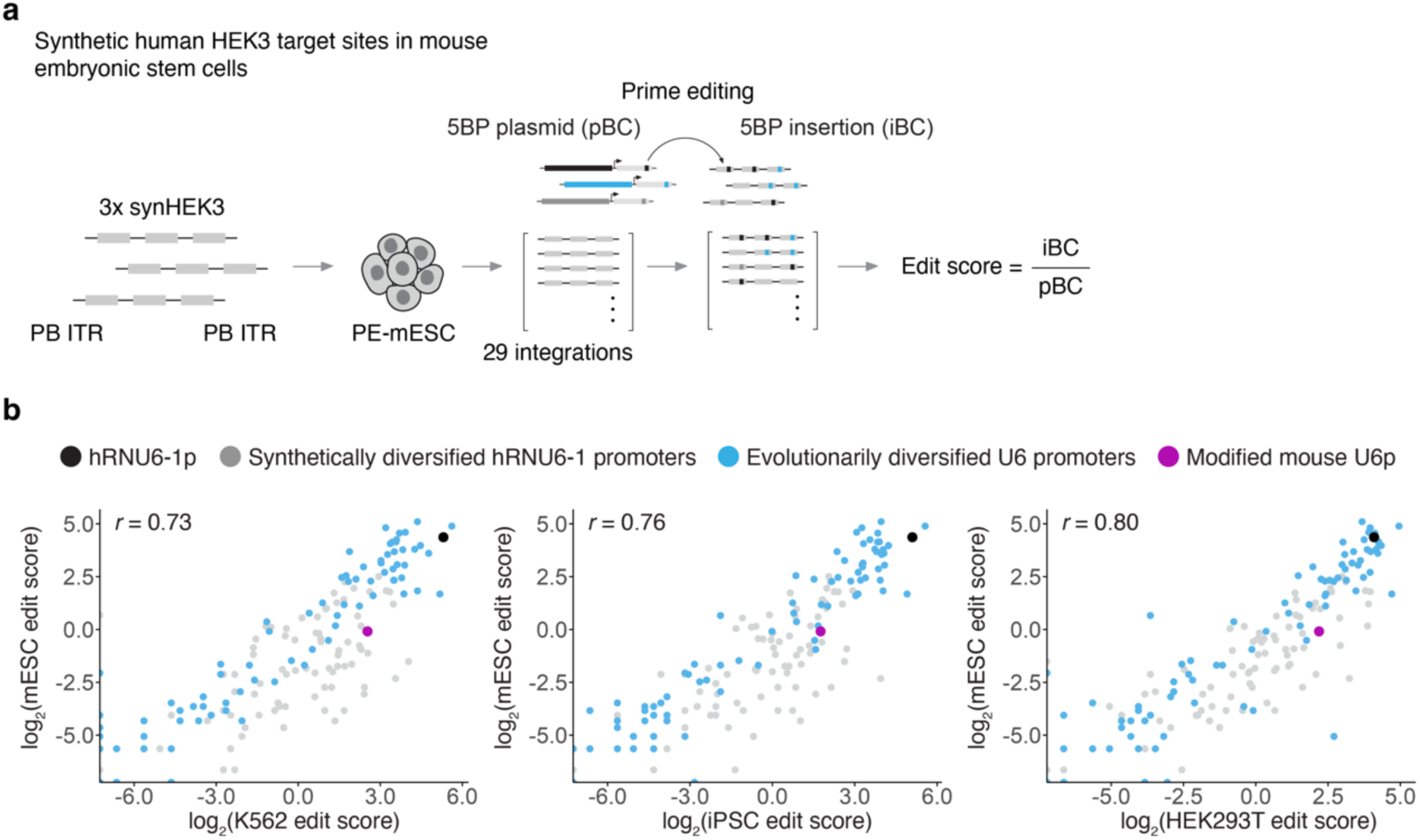
Diversified U6 promoters exhibit consistent activities in human vs. mouse contexts. **a)** PEmax and synthetic human *HEK3* target sites (*synHEK3*) were introduced to mESCs via piggyBac transposition. A monoclonal line with PEmax and an estimated 87 *synHEK3* targets was isolated (29 integrations x 3 *synHEK3* targets per integration). Diversified U6 promoters were assessed for their ability to drive genome editing in mESCs, once again using a multiplex prime editing functional assay. **b)** Pairwise comparison of log-transformed edit scores between three human (x-axes) and one mouse (y-axis) cellular contexts. Pearson correlations, calculated on barcode-normalized edit scores prior to log transformation, are shown.

As in human cells, diversified U6 promoters drove prime editing in mESCs with very strong correlation between technical replicates (*r* > 0.99; **Supplementary Fig. 11**). This higher correlation is probably due to the much larger number of *synHEK3* sites in these engineered mouse cells than endogenous *HEK3* sites in human cell lines (∼87 vs. 2-3), which is expected to decrease measurement noise. Furthermore, results correlated well between human and mouse cells (*r* = 0.73-0.80; **Fig. 4b**; **Table S1**). In mESCs as in human cells, evolutionarily diversified U6 promoters exhibited greater variance in activity (**Fig. 4b**). The human RNU6-1 promoter was again among the top performing promoters in mESCs, consistently outperforming a commonly used, modified mouse U6 promoter^24,50,51^ as well as another mouse U6 promoter that was part of the evolutionarily diversified set (**Fig. 4b**; **Table S1**). Other evolutionarily diversified promoters that were among the most highly active in the human context were similarly highly active in the mouse context (**Fig. 4b**). These results suggest that the widely ranging functional activities of these extant U6 promoters, at least within the species range examined here, are determined by *cis* rather than *trans* differences. As such, they can likely be used across mammalian model systems with the expectation that their activities will be similar to those observed in human cell lines.

### Single-step assembly and deployment of a “ten key” diversified molecular recording array

With functional parts in hand, we sought to test whether these parts were sufficiently sequence-diverse to enable their one-step assembly in yeast, and to then deploy this assembly in mammalian cells. In addition, we sought to assess whether activity measurements for isolated Pol III promoters, scaffolds, and iBCs could be used to predict the activity of novel U6p-pegRNA-iBC combinations, as well as the relative activity of multiple U6p-pegRNA-iBC units assembled into a large array. For this, we designed a ten-unit array of “keys” based on our diversified parts and DNA Typewriter^31^, a temporally resolved molecular recording system that enables sequential insertions of barcodes to a shared DNA Tape (**Fig. 5a**). Sequential records generated with DNA Typewriter can be used to reconstruct various cellular event histories, *e.g.* of cell lineage^31,39^. In designing this diversified molecular recording array, we sought to balance the activity levels of individual U6p-pegRNA-iBC units, as this is expected to yield a greater diversity of sequential editing patterns and thereby maximize the information content of any resulting recordings. Specifically, we paired the top 10 promoters with the top 10 scaffolds in reverse rank order (1st-ranked promoter with 10th-ranked scaffold, 2nd with 9th, etc.; **Fig. 5a**). Further, we paired each U6p-pegRNA unit with specific “NNN<u>GGA</u>” DNA Typewriter barcodes with similar activity levels^31^. A simple multiplicative model of Pol III *x* scaffold *x* iBC edit score predicted that the editing rates of these ten U6p-pegRNA-iBC units would fall within a 2.3-fold range (**Fig. 5a**; **Table S4**).

**Figure 5.**
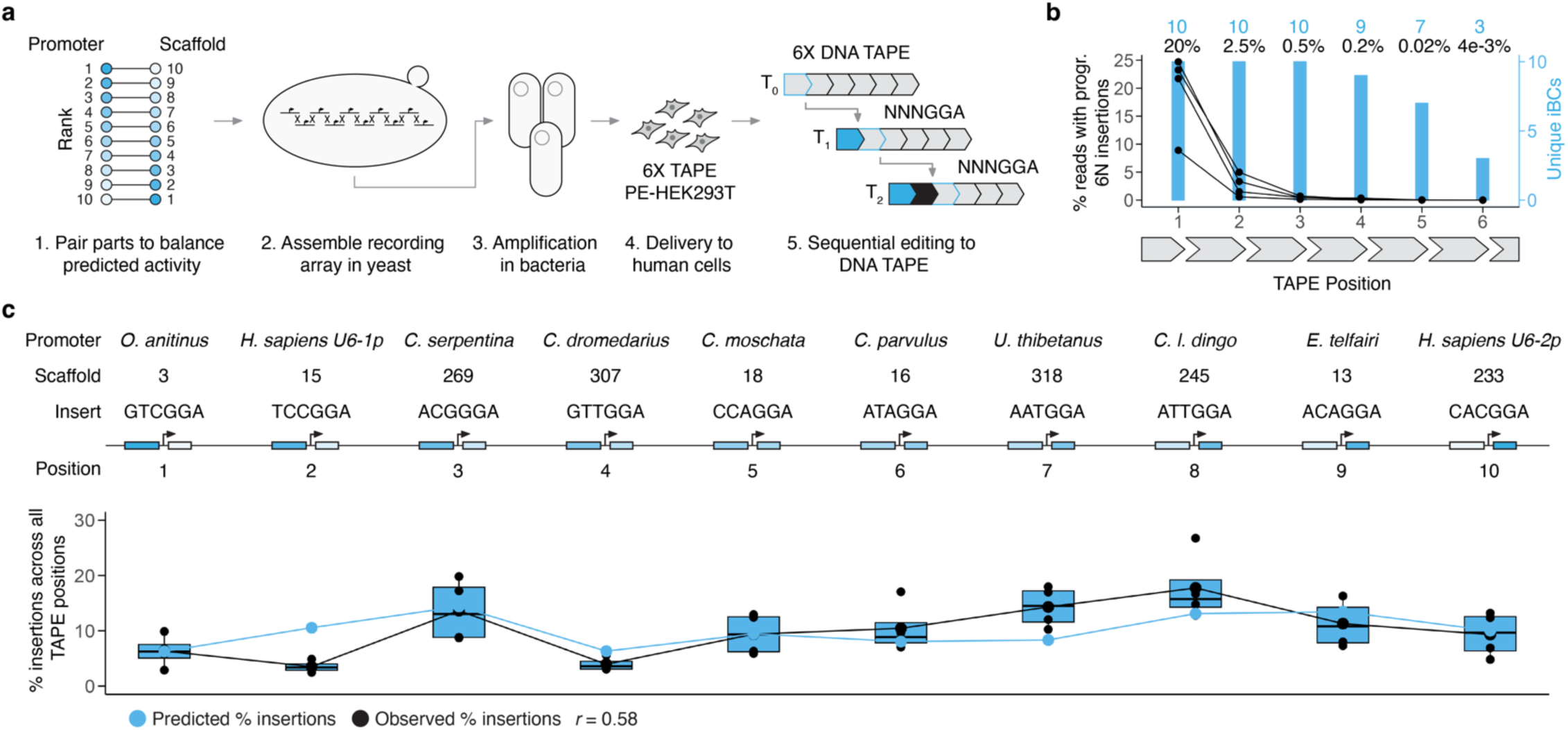
Single-step assembly and functional testing of a “ten key” diversified molecular recording array. **a)** Workflow schematic. 1. Diversified parts are paired in reverse rank order based on individual part activity measurements to balance predicted activity levels. 2. Diversified U6p-pegRNA-iBC units are assembled in one step in yeast. 3. The assembly is recovered, sequence validated and amplified in bacteria. 4. The assembly is delivered to mammalian cells for sequential recording with DNA Typewriter. 5. Each insertion of an NNN iBC and linked GGA key sequence creates a full target site, enabling editing at the next sequential site in DNA Tape. **b)** Editing efficiency (black) and number of unique iBCs recovered (blue) at each of the six sequential sites in DNA Tape. Black dots represent individual transfection replicates (n=4). Higher editing rates in earlier sites are expected due to sequential editing by DNA Typewriter. **c)** Proportion of insertions derived from each of the 10 units in the diversified recording array across all DNA Tape sites. Observed relative editing rates (black) mirror predicted editing rates (blue). Black dots represent individual transfection replicates (n=4). Boxes represent the 25th, 50th, and 75th percentiles. Whiskers extend from hinge to 1.5 times the interquartile range.

We ordered 494-573 bp sequences corresponding to these ten U6p-pegRNA-iBC units flanked by Versatile Genetic Assembly System (VEGAS) adapters^52^ to facilitate their assembly in yeast (**Supplementary Fig. 12**). Additional components of the overall design included piggyBAC inverted terminal repeats (for random integration), *Bxb1* attB sites (for site-specific integration), orthogonal restriction enzymes sites (for isolation of individual units or the entire array), and flanking anti-repressor-elements (for insulation)^53,54^ (**Supplementary Fig. 12**). Following the pooled transformation of 14 fragments to yeast (10 U6p-pegRNA-iBC units, four auxiliary and backbone components), we successfully recovered the complete 15.8 kb ten-unit assembly (**Supplementary Fig. 12**). Whole-construct sequencing revealed only one single nucleotide substitution error that fell at the 5’ end of one of the U6 promoters, upstream of the four core TFBS.

We delivered this “ten key” DNA Typewriter construct to a HEK293T cell line expressing PEmax and multiple integrated copies of a synthetic DNA Tape construct, each with six editable sites for sequential recording (**Fig. 5a**). After 72 hrs, we observed all or a subset of the ten expected NNNGGA barcodes at each of the six sites, at rates that progressively decreased from the first to sixth unit, consistent with sequential editing (**Fig. 5b**). Importantly, we observed insertions corresponding to all 10 U6p-pegRNA-iBC units, and the proportion of edited reads corresponding to each unit was balanced within a few fold at each DNA Tape site where all 10 iBCs were observed (**Fig. 5c**; **Table S4**). Further, the proportion of edited reads predicted by the Pol III *x* scaffold *x* iBC model of our individual part measurements mirrored their observed activities throughout the length of the tandem array, with no obvious systematic bias attributable to the 5’ → 3’ position of the U6p-pegRNA-iBC units (**Fig. 5c**; 2.3-fold predicted range, 4.7-fold observed range; *r* = 0.58; **Table S4**). Taken together, these experiments confirm that our diversified parts are amenable to large-scale assembly in yeast, and that we can predict the activity of novel Pol III promoter-gRNA scaffold-iBC combinations (and tandem arrays thereof) based on the measured activities of individual parts.

### Testing thousands of ancestral, extant, and mutagenized sequences reveals highly active Pol III promoters for mammalian genome editing

Functional candidate parts for genome engineering can be mined from not only extant but also ancestral genomes, *e.g.* as has been done for cytidine deaminases^55^. To further expand the set of sequence- and activity-diversified Pol III promoters available for use in synthetic biology and genome engineering, we leveraged the Zoonomia Project’s 240-species Cactus genome alignment^56–58^ to identify extant and ancestral orthologs of seven Pol III promoters known to be functional in mammalian cells (RNU6-1, RNU6-2, RNU6-7, RNU6-8, RNU6-9, H1, and 7SK promoters). Altogether, we extracted 2,192 unique Pol III promoter sequences, including 1,084 that exactly match at least one extant genome, and 1,108 that solely occur in inferred, ancestral genome(s). We supplemented these mammalian Pol III promoters with saturation mutagenesis libraries encompassing all single nucleotide substitutions and deletions of the human H1 (100 bp, 401 variants including wildtype) and 7SK (243 bp, 973 variants including wildtype) promoters. Altogether, this library contained 3,566 ancestral, extant or mutagenized mammalian Pol III promoters (**Fig. 6a**).

**Figure 6.**
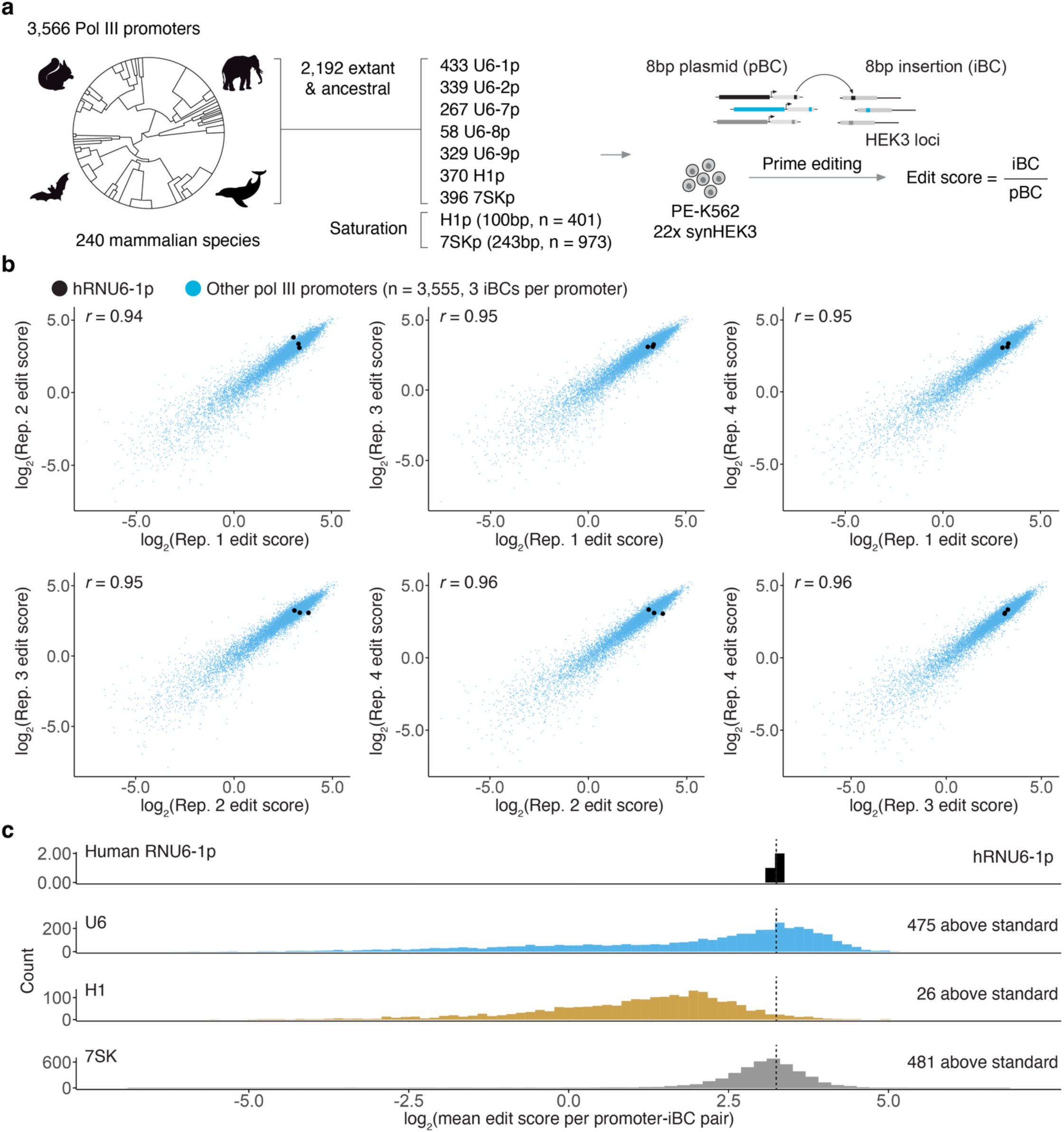
Testing thousands of ancestral, extant, and mutagenized sequences reveals highly active Pol III promoters for genome editing in mammalian cells. **a)** Library design, contents, and multiplex prime editing functional assessment workflow. **b)** Edit scores correlations across the four transfection replicates. Points represent edit scores for the 3 independent iBCs paired with each of the 3,566 promoters (10,698 constructs total). Pearson correlations, calculated on barcode-normalized edit scores prior to log transformation, are shown. **c)** Edit score distributions for the different promoter classes tested in this experiment. The standard human RNU6-1 promoter is shown in the top row, and its mean activity marked with a vertical dashed line.

To facilitate accurate quantification of the relative activities of these promoters, we leveraged insights from earlier experiments. First, given the high technical reproducibility of multiplex prime editing experiments conducted in monoclonal mESCs with large numbers of *synHEK3* target sites (*r* > 0.99; **Supplementary Fig. 11**), we used a monoclonal K562 line with 22 *synHEK3* targets^49^ and PEmax^41^ as our prime editor for these experiments (**Fig. 6a**). Second, we paired each Pol III promoter with 3 independent iBCs (3,566 promoters x 3 iBCs = 10,698 constructs total), accommodating the larger library size by switching from a 5 to 8 bp barcode. To facilitate downstream normalization, we measured the relative insertion activity of all 65,536 possible 8N insertions when driven by the same hRNU6-1p promoter (**Supplementary Fig. 13**).

Following transfection and *synHEK3* amplicon sequencing, we observed the expected insertional edits with strong concordance in edit scores derived from four transfection replicates (*r* > 0.94; **Fig. 6b**). We also observed strong correlation across the three independent iBCs associated with each Pol III promoter *(r* > 0.80; **Supplementary Fig. 14**). This correlation was markedly improved by correcting for relative barcode insertion efficiency (r = 0.48-0.51 before vs. 0.80-0.81 after barcode correction; **Supplementary Fig. 14**). This result reinforces the importance of having relative activity measurements for all iBCs used, particularly for longer iBCs, which exerted greater influence on raw edit scores than shorter barcodes (**Supplementary Fig. 2**; **Supplementary Fig. 13**).

Global analyses of this screen revealed a broad range of mammalian Pol III promoter activity levels, with clear differences between the activity distributions of the classes of elements tested. Evolutionary orthologs of the H1 promoter exhibited weaker activity than orthologs of U6 or 7SK promoters (**Fig. 6c**), consistent with our earlier comparisons of the short H1 and miniaturized U6 promoters vs. the full length U6 promoter (**Fig. 3a**). Also consistent with expectation, saturation mutagenesis of the human H1 and 7SK promoters highlighted the four core TFBS as particularly constrained, while also identifying numerous tolerated and activity-enhancing SNVs which could be leveraged for additional diversification (**Supplementary Fig. 15**). Interestingly, as compared with U6, the H1 and 7SK Pol III promoters were much more tolerant of single nucleotide deletions in their TATA boxes, but much less tolerant of mutations in the SPH or PSE elements (**Fig. 3b**; **Supplementary Fig. 15**).

As in earlier screens, hRNU6-1p was among the most highly active promoters (**Fig. 6c**). Remarkably however, we also identified 982 promoters that outperformed hRNU6-1p across all iBCs (982/3566 or 28%, including 475 U6, 26 H1, and 481 7SK promoter orthologs; median 1.3-fold increase over hRNU6-1p; **Fig. 6c**; **Table S5**). 408/982 (42%) of these hRNU6-1p outperformers were not present in any extant mammalian genome in the Zoomania Project, highlighting the potential value of inferred, ancestral genome(s) as a source of noncoding regulatory parts for synthetic biology. These included the most active Pol III promoter in this experiment, a 7SK promoter ortholog from an intermediate ancestral rodent genome that drove prime editing at *synHEK3* sites with 2.6-fold greater activity than hRNU6-1p. Other top-performers derived from saturation mutagenesis (25%) or extant genomes (33%), the latter including Pol III promoters from the genomes of the java mouse deer (*Tragulus javanicus*), long-tongued fruit bat (*Macroglossus sobrinus*), Linnaeus’s two-toed sloth (*Choloepus didactylus*), and one of our closest relatives, the bonobo (*Pan paniscus*) (**Table S5**).

We suspect that the much higher proportion of Pol III promoters whose activities exceed hRNU6-1p in this screen as compared with the earlier screen follows from sampling an order of magnitude more sequences from more closely related species, with less attention to ensuring their sequence divergence. Nonetheless, this set is sufficiently large so as to enable the selection of subsets that are highly sequence-diverse so as to facilitate yeast-based assembly. For example, of the 481,687 possible pairwise comparisons among the 982 Pol III promoters that outperformed hRNU6-1p, there exist subsets of at least 205 that satisfy *L_max_* < 40 (**Supplementary Fig. 16**). This effectively provides a large set of yeast-assembly-compatible Pol III promoters that are as or more active than hRNU6-1p for driving genome editing.

While our main goal was to generate diversified parts to facilitate genome engineering, synthetic biology, and molecular recording, this experiment incidentally mapped the functional landscape of mammalian Pol III promoter evolution. Superimposing Pol III promoter activity levels onto a phylogenetic tree revealed clear trajectories of Pol III promoter evolution (**Supplementary Fig. 17**). For example, hRNU6-1p orthologs display strong, deeply conserved activity across extant and reconstructed ancestral species (**Fig. 7a**; **Supplementary Fig. 17**). In contrast, hRNU6-9p orthologs derived from most species exhibit low activity, but appear to have acquired strong activity in the primate lineage specifically, including humans (**Fig. 7b**).

**Figure 7.**
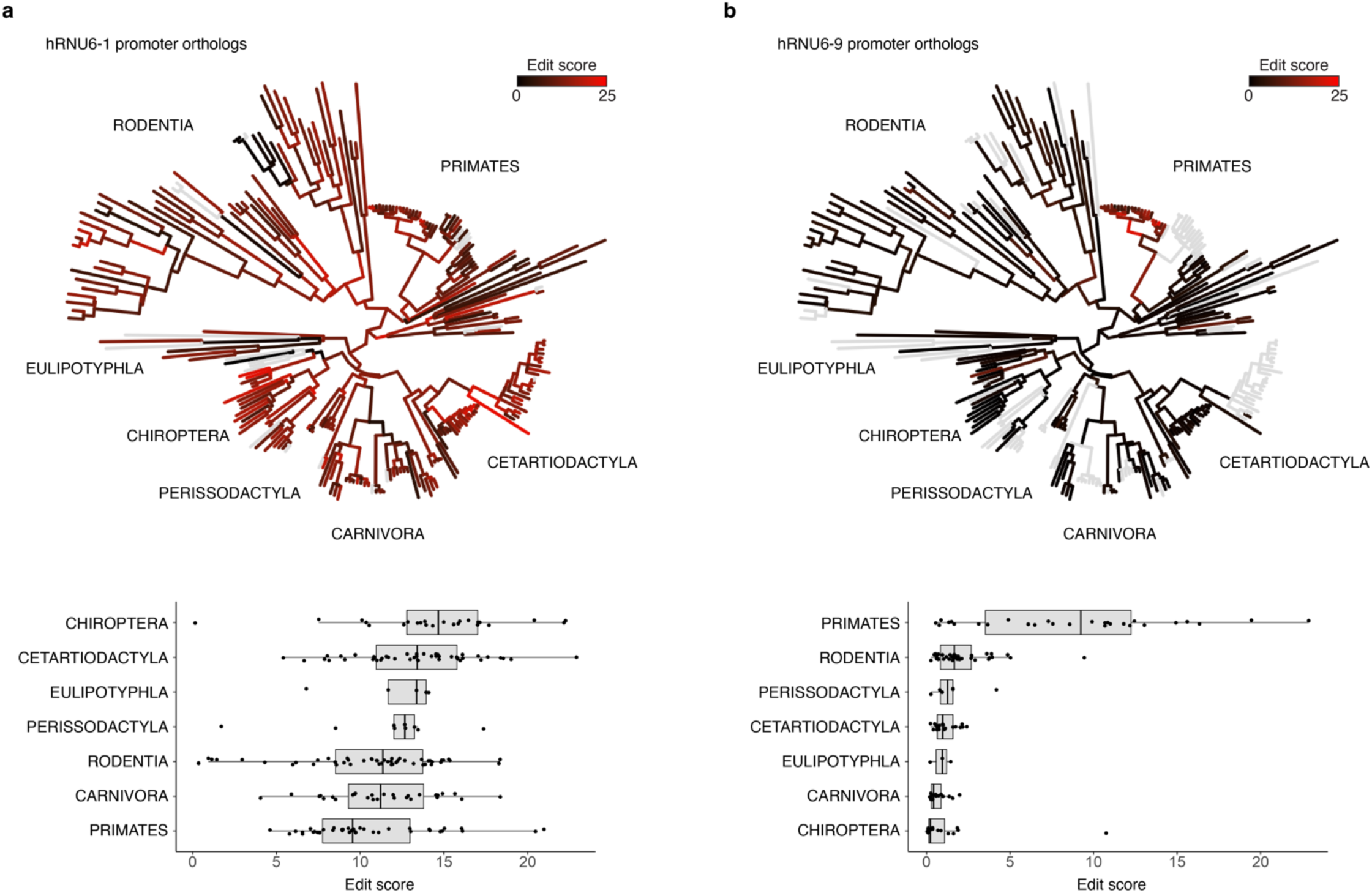
Functional trajectories of mammalian Pol III promoter evolution. **a)** (top) Phylogenetic tree depicting the evolutionary relationships and edit scores of extant and ancestral hRNU6-1p orthologs. (bottom) Comparison of edit scores for hRNU6-1p orthologs from major orders. hRNU6-1p orthologs display strong, deeply conserved activity across extant and reconstructed ancestral species. Boxes represent the 25th, 50th, and 75th percentiles. Whiskers extend from hinge to 1.5 times the interquartile range. **b)** Same as panel **a** but for extant and ancestral hRNU6-9p orthologs.

## Discussion

Here we report sequence-diversified and miniaturized parts for multiplex CRISPR-based genome engineering in mammalian cells. These parts exhibit consistent performance across multiple cell contexts, including the workhorses of functional genomics technology development (HEK293T, K562) and the starting points for diverse organoid and *in vivo* models (human iPSCs, mouse ESCs). Parts in each class (Pol III promoters, guide RNA scaffolds) exhibit reproducible activity spanning over three orders of magnitude (and applied together potentially over six orders of magnitude). Many of these parts outperform the widely used standard parts, and may be useful simply for maximizing genome editing rates in routine experiments.

More sophisticated applications may include any genome engineering or synthetic biology project in which simplified assembly, miniaturization and/or activity titration would be beneficial. Although we focused on simplified assembly for molecular recording in the follow-up experiments reported here, other applications which will benefit from both simplified assembly and miniaturized parts include packaging multiple U6p-gRNA cassettes into recombination-prone viral vectors commonly used in CRISPR screens^24,34,59–62^ or for gene therapy, while applications which will benefit from activity titration include the design and implementation of complex genetic circuits. Although genome editing activity can also be titrated via spacer mismatches as demonstrated by Jost *et al.*^42^, titrating activity via the Pol III promoter or gRNA scaffold has the advantage of being generic across targets. Finally, we note that our Pol III parts list may also be useful for non-CRISPR synthetic biology applications relying on quantitative control of short RNA expression.

The diversity of activity levels of evolutionarily diversified Pol III promoters is interesting in its own right, and appears to be a consequence of divergence in *cis* rather than *trans* regulation. The evidence for this is that U6 promoters from diverse vertebrate species exhibited consistent activities in multiple human and mouse cellular contexts. For example, human and platypus U6 promoters outperformed multiple mouse U6 promoters in all human and mouse cell lines tested. This is not terribly surprising, as we presume that the *trans* acting factors underlying Pol III transcription are deeply conserved, much more so than the sequences of individual Pol III promoters, analogous to what is clearly the case for the much more deeply studied Pol II transcription factors and Pol II enhancers/promoters at which they act. The beneficial implication is that the activity levels documented here for individual parts are likely to be consistent in untested contexts (*e.g.* other human or mouse cell types; other mammalian species).

We based our multiplex functional assay on prime editing because this permitted the use of part-specific insertional barcodes, facilitating straightforward quantitation of the relative activity of thousands of parts in a single experiment. Quantifying genome editing rather than RNA abundance was critical, as diversified Pol III promoters can have variable amounts of Pol II activity, which can produce alternative transcripts that are abundant yet fail to drive genome editing^39,48,63,64^. We elected to target *HEK3* because of its well-documented efficiency for insertional prime editing^38,31,39^. However, we expect the activities estimated here to generalize not only across cellular/species contexts, but also across target sites. Indeed, the standard U6 promoter and gRNA/pegRNA scaffold have been successfully used to target countless loci, and when quantified the effects of spacer-scaffold interactions are minor^27,46,65^.

For similar reasons, we predict that diversified scaffolds will also be combinable with other variations on Cas9-mediated genome editing both at the protein (*e.g.* nuclease editing, CRISPR i/a editing, etc.) and guide (*e.g.* epegRNAs with structured motifs at their 3’ end^46^) levels. Indeed, the A-U flip design has recently been used successfully with epegRNAs for improved performance^46^, and we expect the same will be possible with other high-performance scaffold alternatives identified here. Further, the advent of PE7, which fuses an endogenous human RNA-binding domain to PEmax, offers performance on par with epegRNAs while enabling use of shorter, less repetitive standard pegRNAs such as the ones diversified here (probably by conferring pegRNA stability through protein-binding rather than secondary structure)^66^. Similarly, the parts described here may be synergistic with Cas12a arrays and related approaches to multiplex gRNAs in a single transcript, *e.g.* by enabling ‘nested multiplexing’ through assembly and delivery of multiple independent gRNA arrays on a single construct with multiple diversified and/or miniaturized U6 promoters^67–69^.

In our view, among the most exciting use-cases for this parts list lie in the field of molecular recording^4,8,70^. Following up on with our goals in setting out in this direction, we demonstrated that these sequence-diversified parts are amenable to single step assembly in yeast and deployment in mammalian cells as a single-locus, ten-key DNA Typewriter. Further, these experiments revealed that the activity of novel Pol III promoter-pegRNA-iBC combinations (and arrays thereof) can be predicted based on individual part activity measurements, something that was unclear at the outset of this work. Using these parts and following the strategy we have demonstrated here, one could imagine assembling, in yeast and as a single locus, many more iterations and combinations of multi-unit arrayed CRISPR clocks^71^, transcriptional^39^ or lineage recorders^20,23,31,72,73^ that write to their DNA recording medium at different rates in parallel, to concurrently access different temporal resolutions and time scales.

Zooming out, we envision that the strategy taken here, namely combining evolutionary mining with rational design and multiplex functional assays, will advance the realization of a long-standing goal of synthetic biology --the delineation of sequence-diversified, functionally-diversified, cross-compatible “parts”, that can be routinely and cost-effectively assembled to build complex genetic circuits that will behave in a predictable manner^9,10^. A further vision is that the quantitative characterization of these parts will essentially serve as “pre-training” for generative models that can *de novo* design circuits that function as predicted and allow us to access a vast range of possibilities within the space of intracellular circuits.

## Supplementary figures

**Figure S1.**
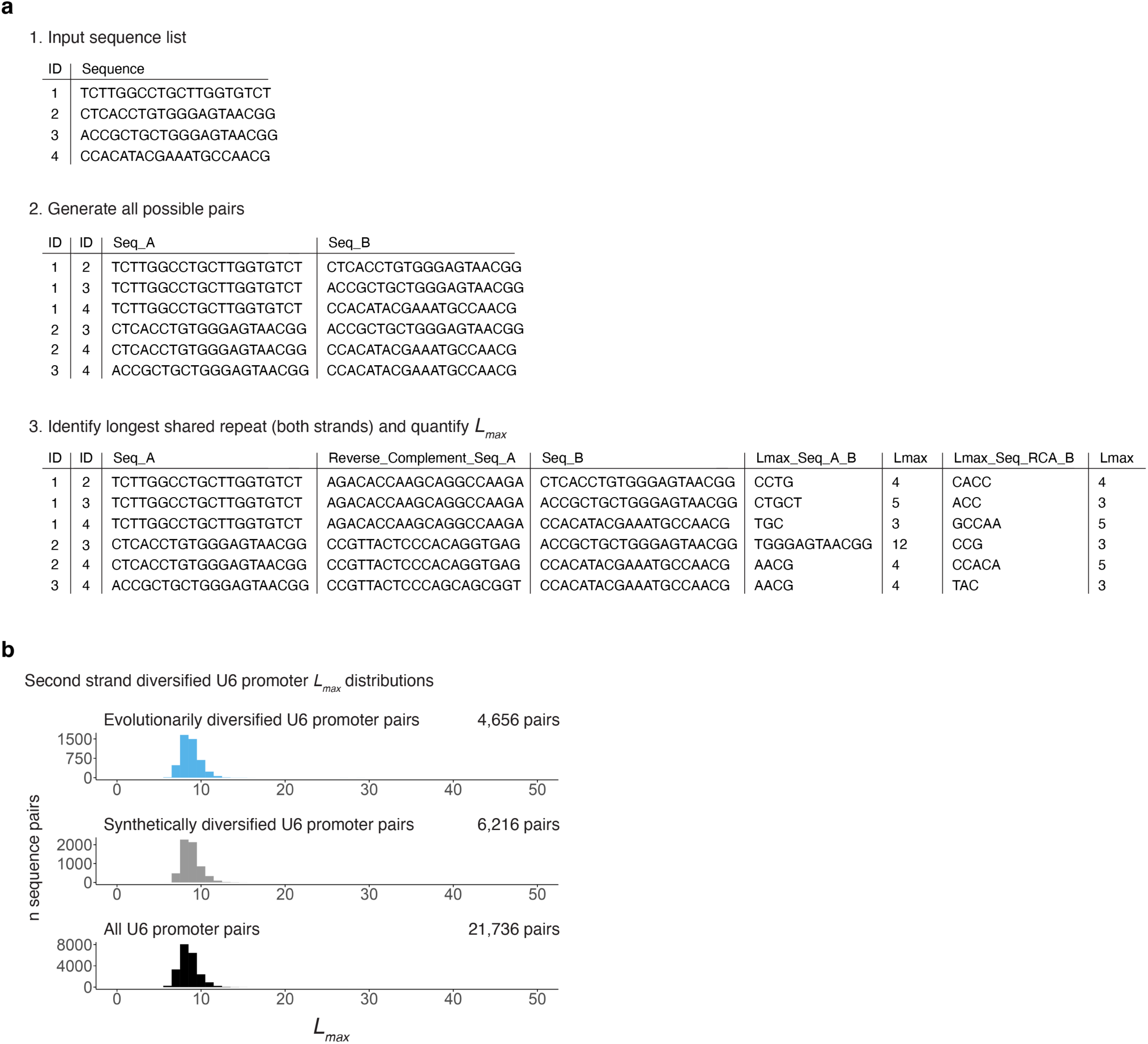
*L_max_* calculation example and second strand diversified U6 promoter library *L_max_* distributions. **a)** Illustrative example of L*_max_* calculations using a set of four 4 gRNA sequences (gRNAs are unrelated to the present work). A set of sequences is input, all possible pairs are generated, the longest shared repeat between each pair is identified (considering both strands), and *L_max_* is reported. **b)** *L_max_* distributions quantifying the maximal shared repeated length between all possible pairs of sequences for the diverse species U6 promoter library (n=97; 4,656 pairs), synthetic hRNU6-1p library (n=112; 6,216 pairs) and combined set (n=209; 21,736 pairs) for reverse complement comparisons. See **Supplementary Fig. 1b** for *L_max_* distributions for same strand comparisons. Note that in practice the longest shared repeat among sequence pairs in the present libraries were overwhelmingly between pairs in the same orientation due to shared functional elements.

**Figure S2.**
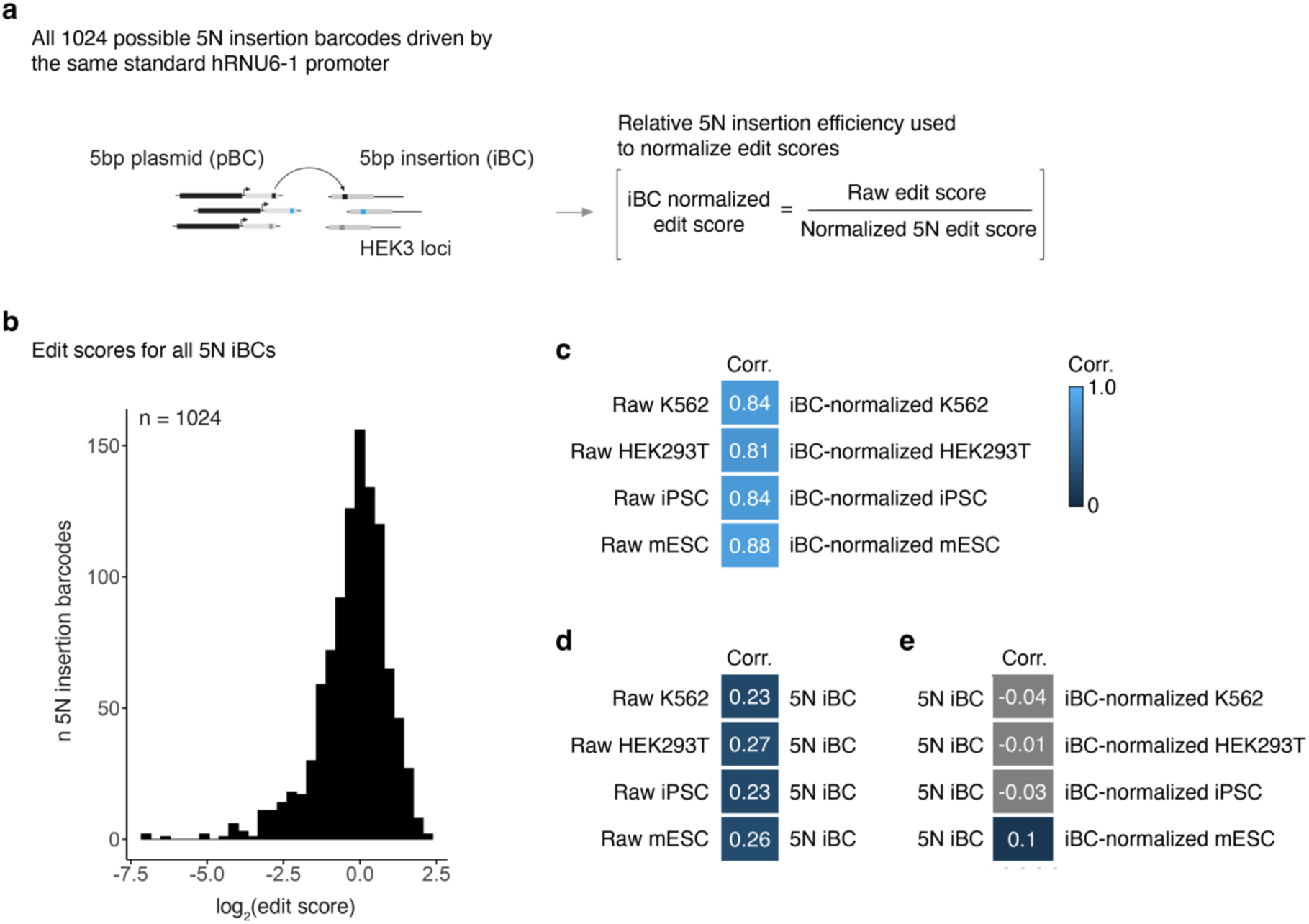
Relative edit scores of all possible 5N insertion barcodes and strategy for edit score normalization. **a)** A library of pegRNAs programmed to insert all 1024 possible 5N iBCs was driven by the standard human RNU6-1 promoter to assess their relative insertion efficiencies. The resulting iBC edit scores, calculated by dividing each 5 bp sequence’s insertion frequency at *HEK3* by its frequency in the pegRNA library, was used to normalize raw edit scores for diversified U6 promoters paired with a given 5N iBC. **b)** Distribution of edit scores for all 5N iBCs driven by the standard human RNU6-1 promoter. Edit scores for 5N iBCs were generally very similar (all 1024 barcodes drove detectable editing, and 905/1024 (88%) fell within 3-fold of the median score). **c)** Correlation between raw edit scores for diversified U6 promoters paired with different 5N iBCs and iBC-normalized edit scores. **d)** Correlation between raw edit edit scores and relative 5N insertion efficiencies. **e)** Correlation between relative 5N iBC efficiencies and iBC-normalized edit scores for diversified U6 promoters. Accounting for relative 5N iBC insertion efficiency effectively corrected for their relatively minor influence on diversified U6 promoter edit scores.

**Figure S3.**
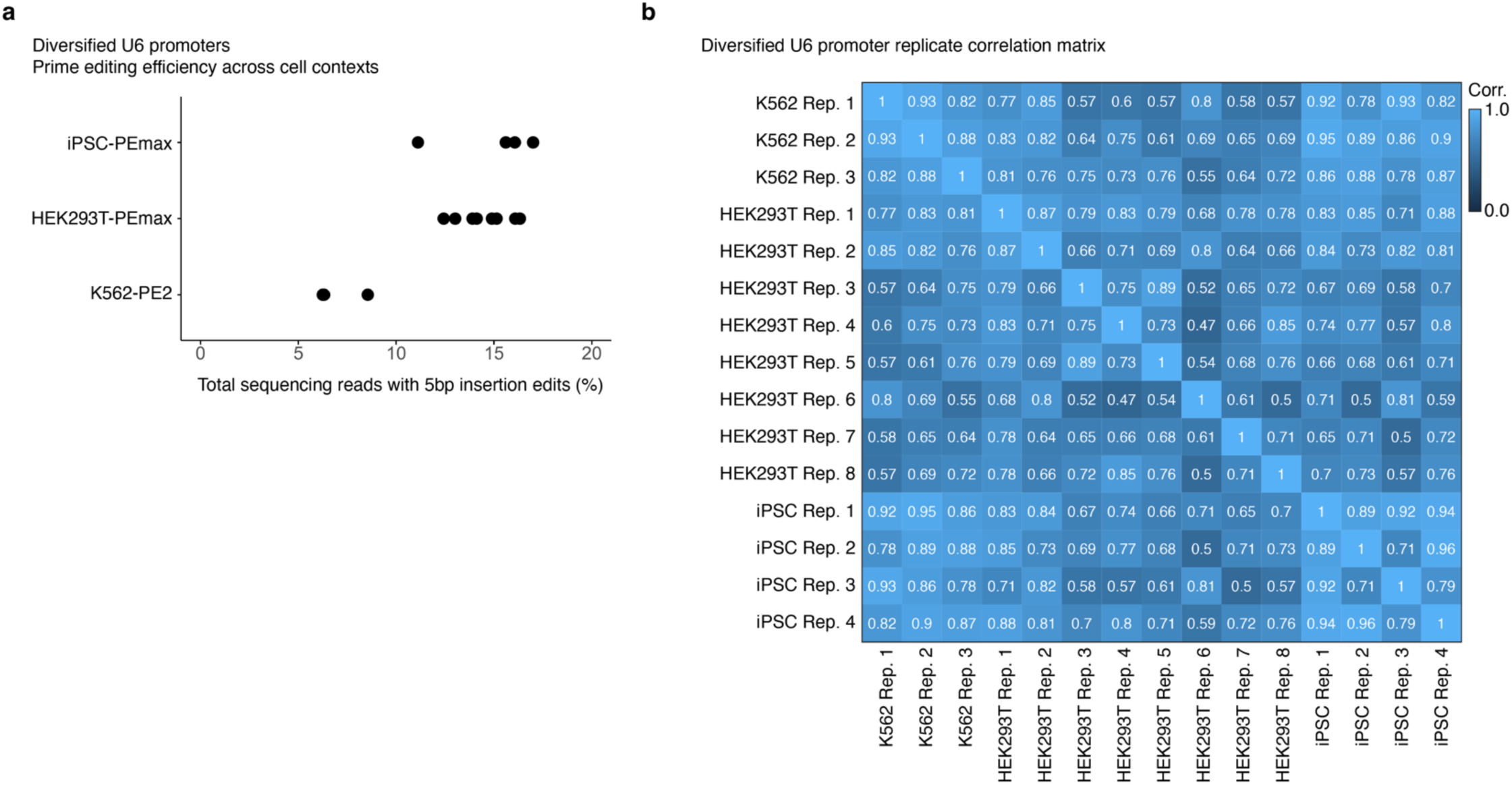
U6 promoter diversification experiment: prime editing efficiencies and edit score replicate correlations across cellular contexts. **a)** Prime editing efficiency of the diversified U6 promoter library across human cellular contexts. Cell lines expressing an optimized PEmax construct^41^ displayed higher editing scores than the K562 line expressing the original PE2 construct^38^, as expected. **b)** Correlation of diversified U6 promoter edit scores across cellular contexts for individual transfection replicates.

**Figure S4.**
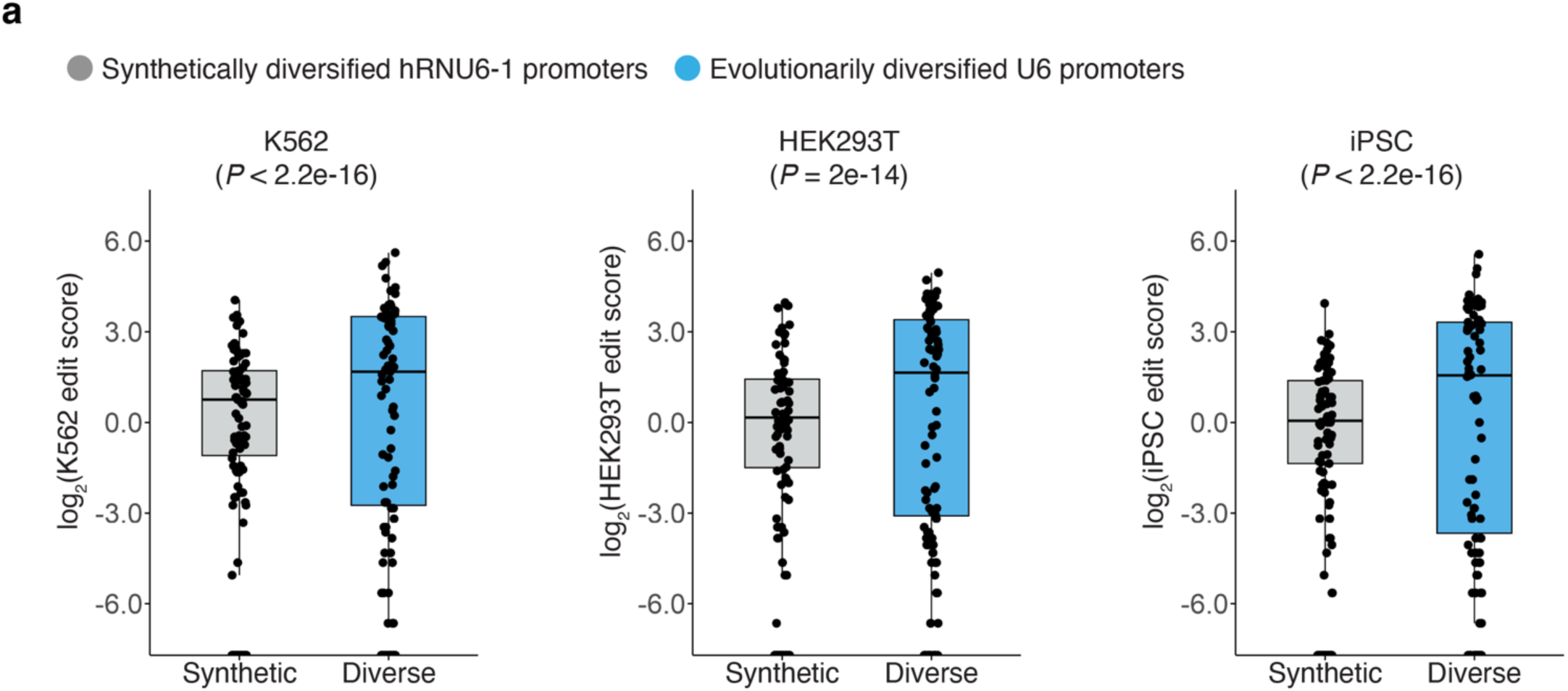
Comparison of synthetically and evolutionarily diversified U6 promoters. **a)** Comparison of edit score distributions for synthetically (gray) and evolutionarily (blue) diversified U6 promoters in three human cellular contexts. Evolutionarily diversified U6 promoters exhibited a greater diversity of activity levels. *P*-values from an *F*-test for equality of variance are shown above. Boxes represent the 25th, 50th, and 75th percentiles. Whiskers extend from hinge to 1.5 times the interquartile range.

**Figure S5.**
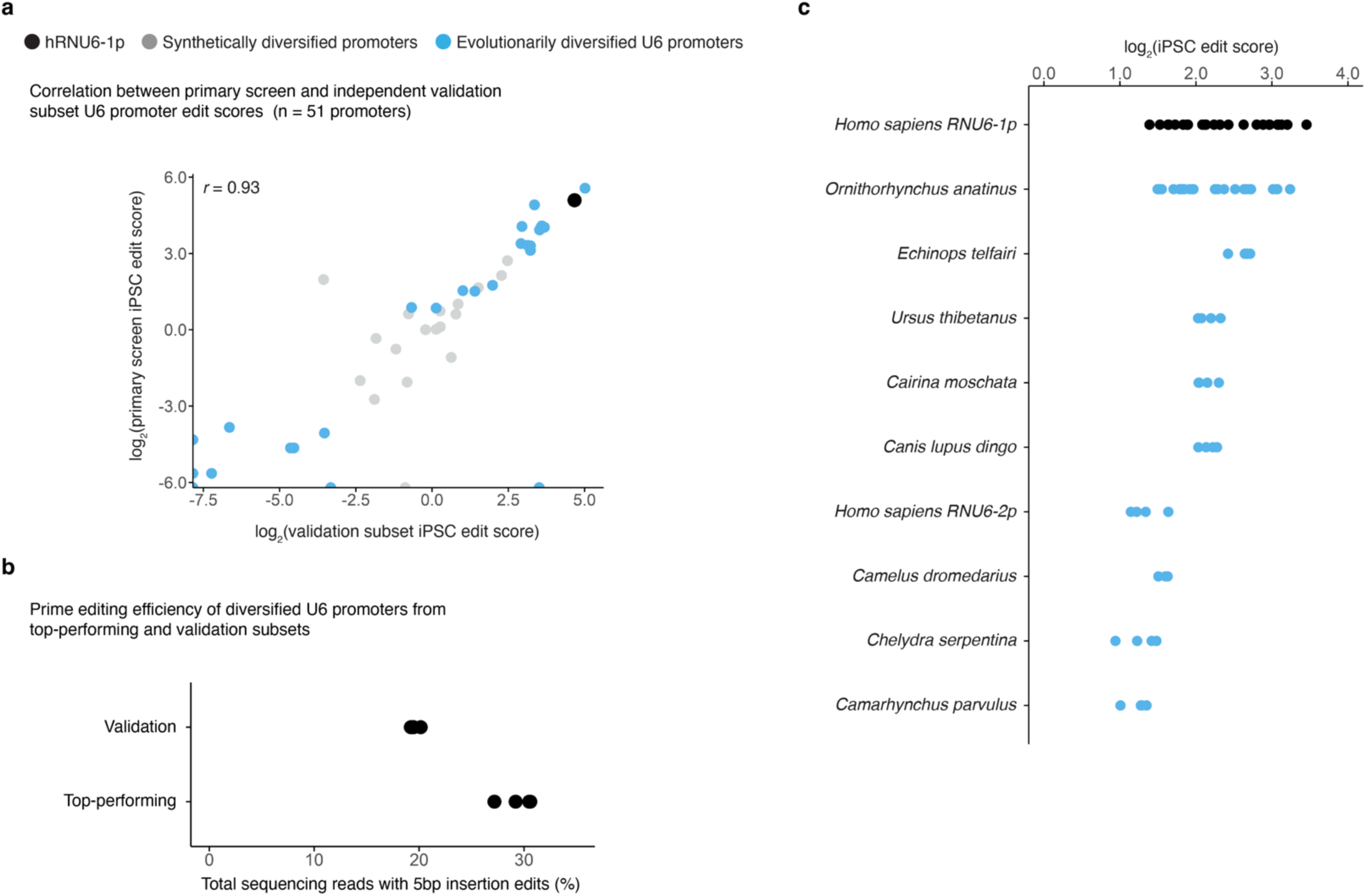
Independent validation of diversified U6 promoters. **a)** Correlation between edit scores in PEmax-iPSCs for diversified U6 promoters in the primary screen vs. an independently cloned and tested subset of 51 diversified promoters representing a range of activity levels. Results correlated strongly between the primary screen and validation experiment. Pearson correlations, calculated on barcode-normalized edit scores prior to log transformation, are shown. **b)** Prime editing efficiencies of the validation subset and top-performing diversified promoters libraries in PEmax-iPSCs, for transfections conducted in quadruplicate. **c)** Edit scores of the top-performing diversified U6 promoters each tested with alternate iBCs that differ from the iBC used in the primary screen in quadruplicate. The standard human RNU6-1p and *Ornithorhynchus anatinus* U6 promoters were each tested with six iBCs in quadruplicate. All top-performing promoters shown here drove robust editing when tested with alternate iBCs and had similar edit scores (within 2.6-fold of standard human RNU6-1p).

**Figure S6.**
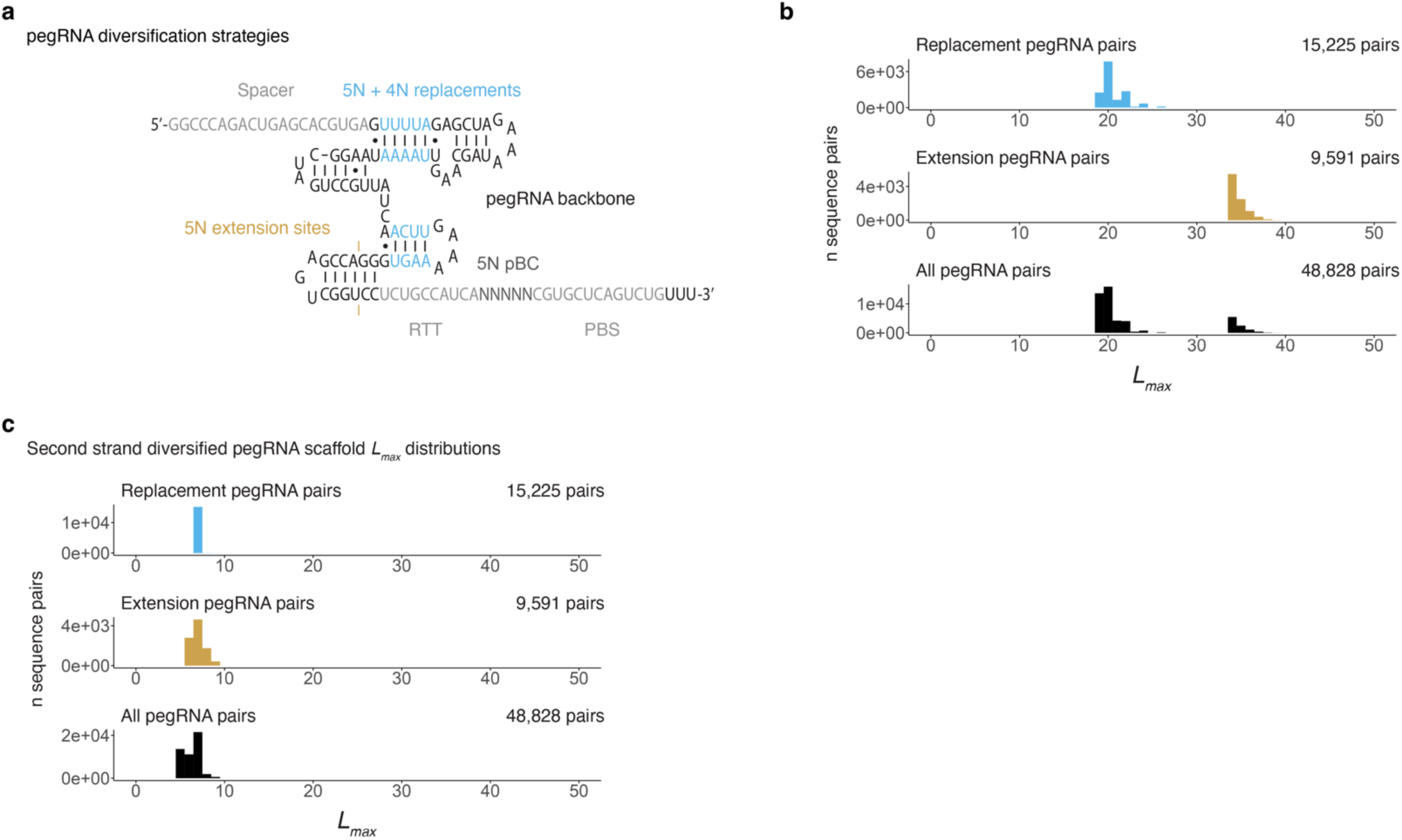
Diversified pegRNA library design and *L_max_* distributions. **a)** Predicted structure of a *HEK3* targeting pegRNA with a standard scaffold sequence programmed to insert a 5N iBC at the *HEK3* locus. Sites where random, complementary 5N and 4N replacements (blue) and extensions (orange) were introduced to diversify scaffold sequences are shown. **b)** *L_max_* distributions quantifying the maximal shared repeated length between all possible pairs of sequences for the replacement scaffolds (n=174, 15,225 pairs), and extension scaffolds (n=138; 9,591 pairs) and combined set including the standard sequence (n=313, 48,828 pairs), in the same orientation. **c)** *L_max_* distributions as in panel **b** but for reverse complement comparisons.

**Figure S7.**
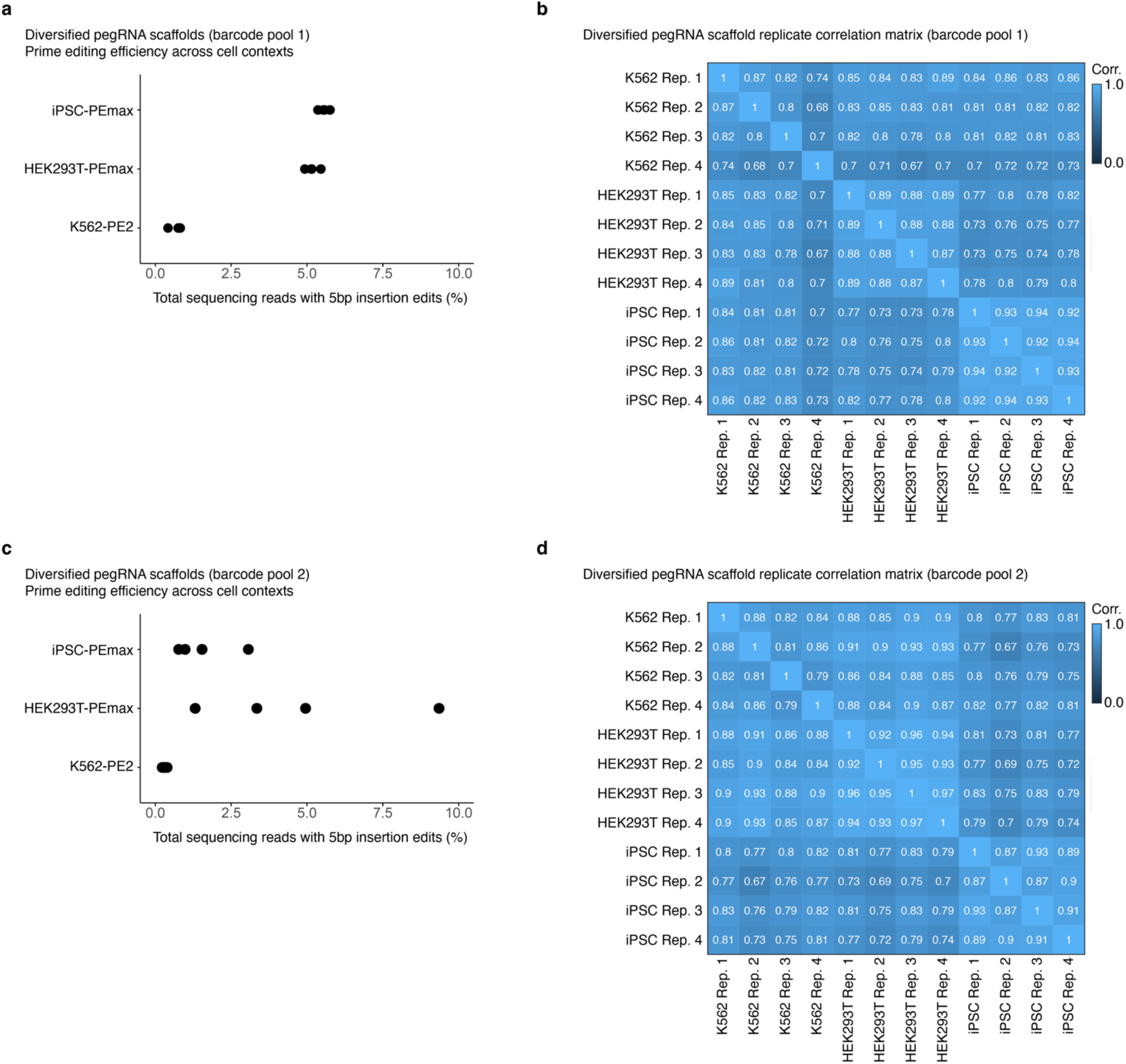
Scaffold diversification experiment: prime editing efficiencies and edit score replicate correlations across cellular contexts. **a)** Prime editing efficiencies of the diversified pegRNA scaffold library across cellular contexts for the first independent barcode pool. Cell lines expressing an optimized PEmax construct^41^ displayed higher editing scores than the K562 line expressing the original PE2 construct^38^, as expected. **b)** Correlation of diversified pegRNA scaffold edit scores across cell contexts for individual transfection replicates of the first independent barcode pool. Pearson correlations, calculated on barcode-normalized edit scores prior to log transformation, are shown. **c)** Same as panel **a** but for the second independent barcode pool. **d)** Same as panel **b** but for the second independent barcode pool.

**Figure S8.**
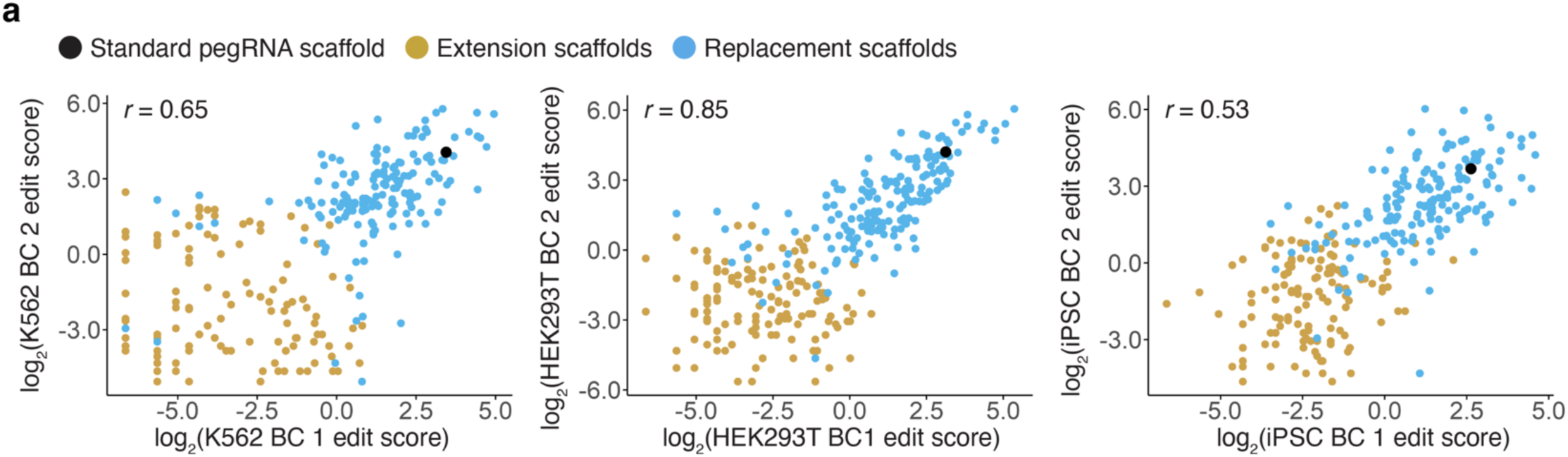
Diversified pegRNA edit score correlations between independent barcode pools. **a)** Correlation of diversified pegRNA scaffold edit scores generated from two independent barcode pools, in each of three human cellular contexts. Pearson correlations, calculated on barcode-normalized edit scores prior to log transformation, are shown.

**Figure S9.**
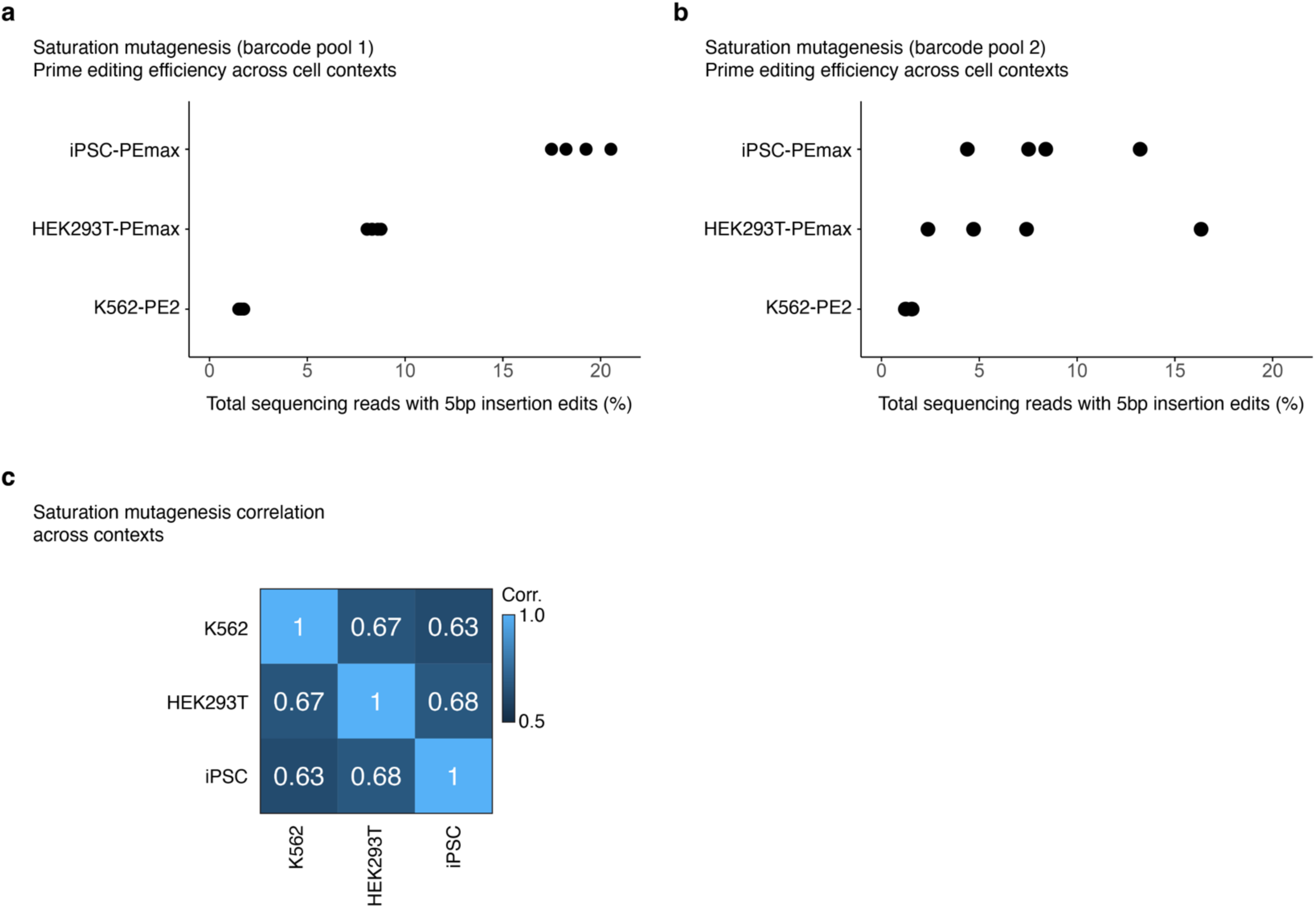
Saturation mutagenesis experiment: prime editing efficiencies and edit score correlations across cellular contexts. **a)** Prime editing efficiencies of the miniaturized hRNU6-1p-pegRNA cassette saturation mutagenesis library across cellular contexts for the first independent barcode pool. Cell lines expressing an optimized PEmax construct^41^ displayed higher editing scores than the K562 line expressing the original PE2 construct^38^, as expected. **b)** Same as panel **a** but for the second independent barcode pool. **c)** Correlation of saturation mutagenesis variant edit scores across cellular contexts. Pearson correlations based on edit scores from both barcodes prior to log transformation, are shown.

**Figure S10.**
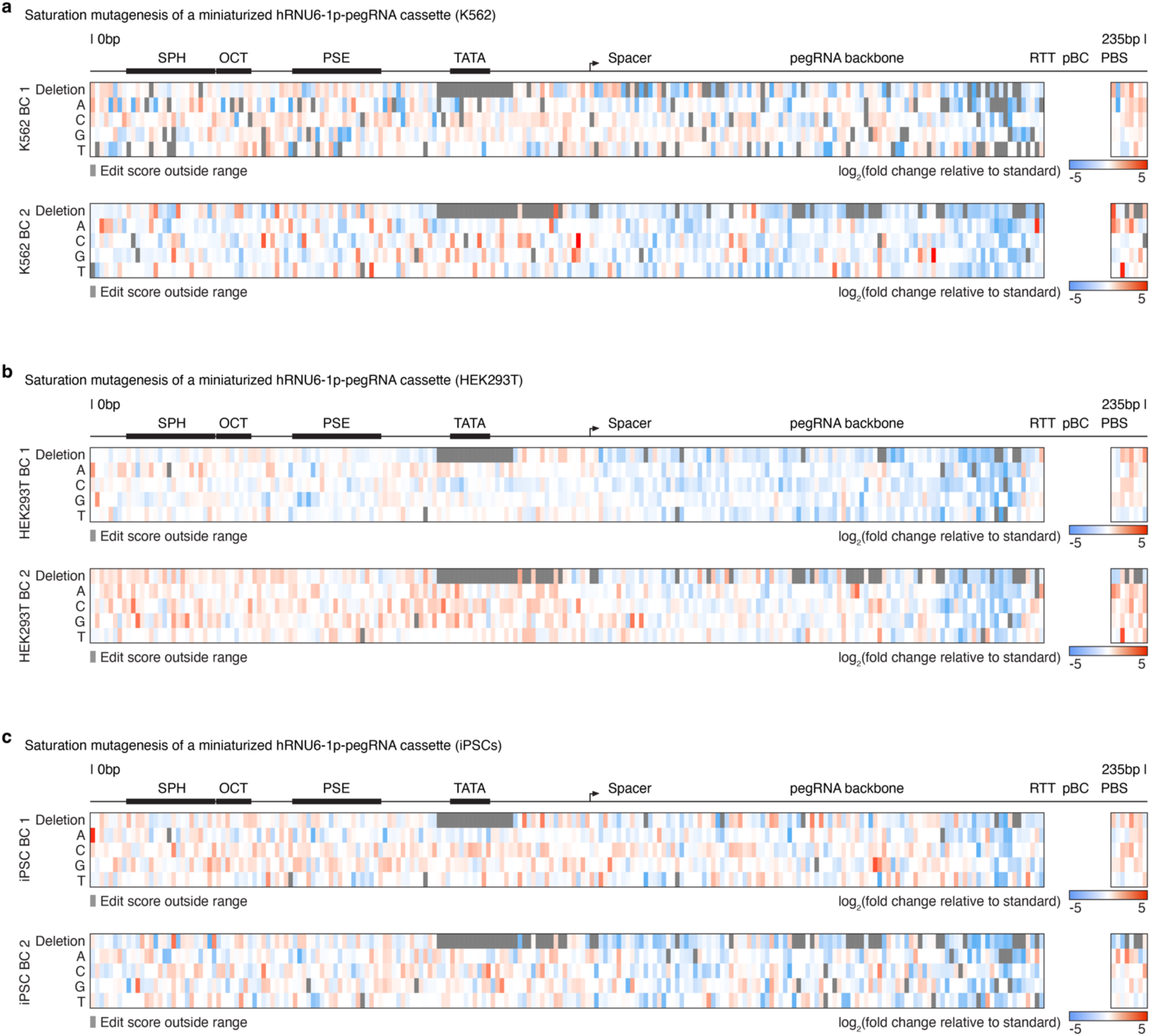
Saturation mutagenesis variant effect maps of miniaturized hRNU6-1p-pegRNA cassettes across cellular contexts. **a)** Saturation mutagenesis variant effect maps of a miniaturized hRNU6-1p-pegRNA cassette tested in K562 cells. Log-transformed fold-change in edit scores relative to the wildtype cassette for the first (top) and second (bottom) independent barcode pools are shown. **b)** Same as panel **b** but experiment conducted in HEK293T cells. **c)** Same as panel **b** but experiment conducted in iPSCs. Edit scores were not calculated for the unboxed region surrounding the pBC, as exact matches spanning this region were required for edit quantification.

**Figure S11.**
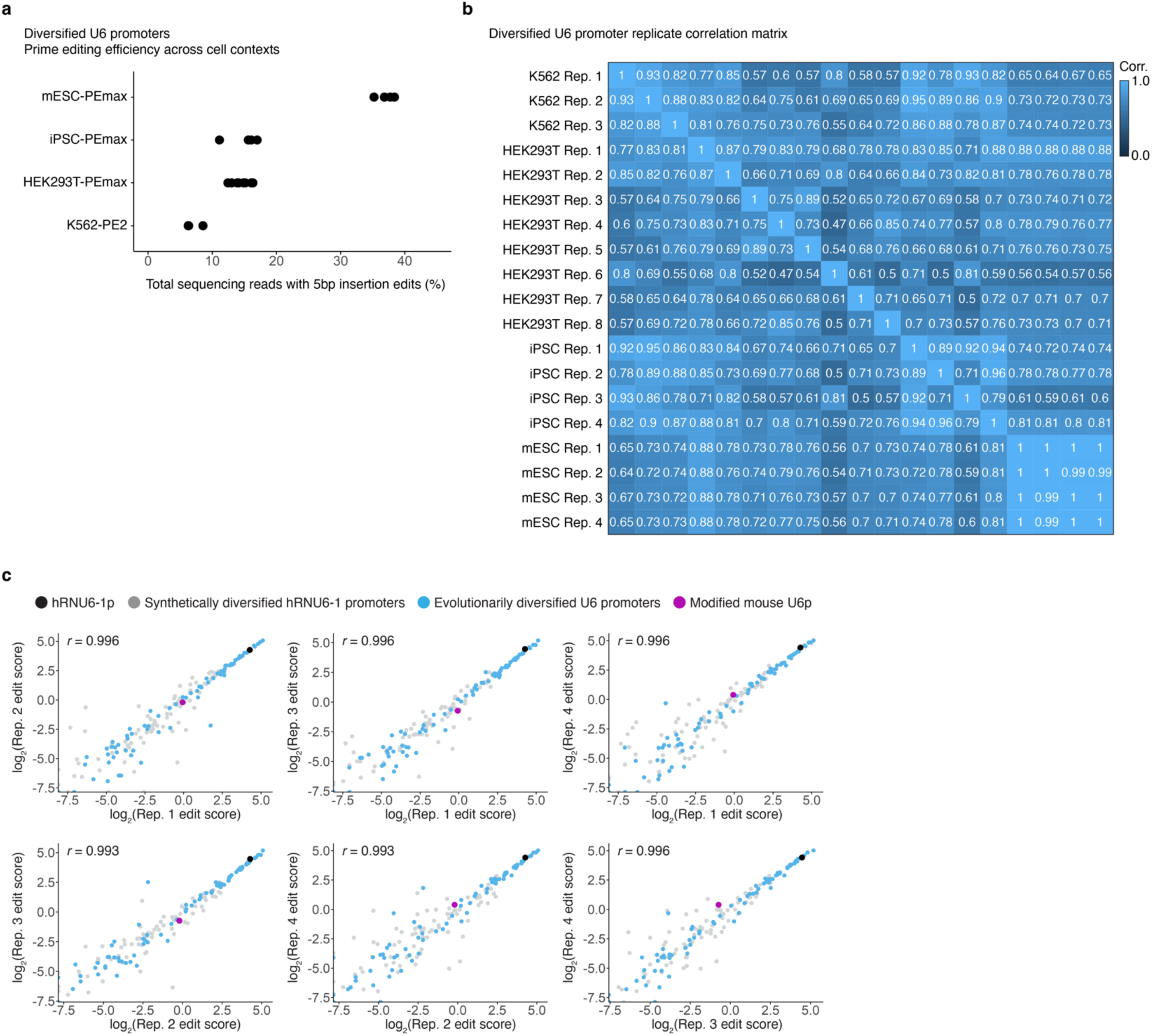
Extension of the U6 promoter diversification experiment to mouse embryonic stem cells: prime editing efficiencies and edit score replicate correlations. **a)** Prime editing efficiency of the diversified U6 promoter library in mESCs vs. human cell contexts. The editing efficiency was considerably higher in the mESC line, possibly because it was engineered to harbor many *synHEK3* target sites per cell, or possibly for other reasons. **b)** Correlation of diversified U6 promoter edit scores across cell contexts for individual transfection replicates. Pearson correlations, calculated on barcode-normalized edit scores prior to log transformation, are shown. Note certain mESC technical replicates had a Pearson correlation above 0.995 and round to 1 in the visualization. **c)** Edit scores correlations across the four transfection replicates. Pearson correlations, calculated on barcode-normalized edit scores prior to log transformation, are shown.

**Figure S12.**
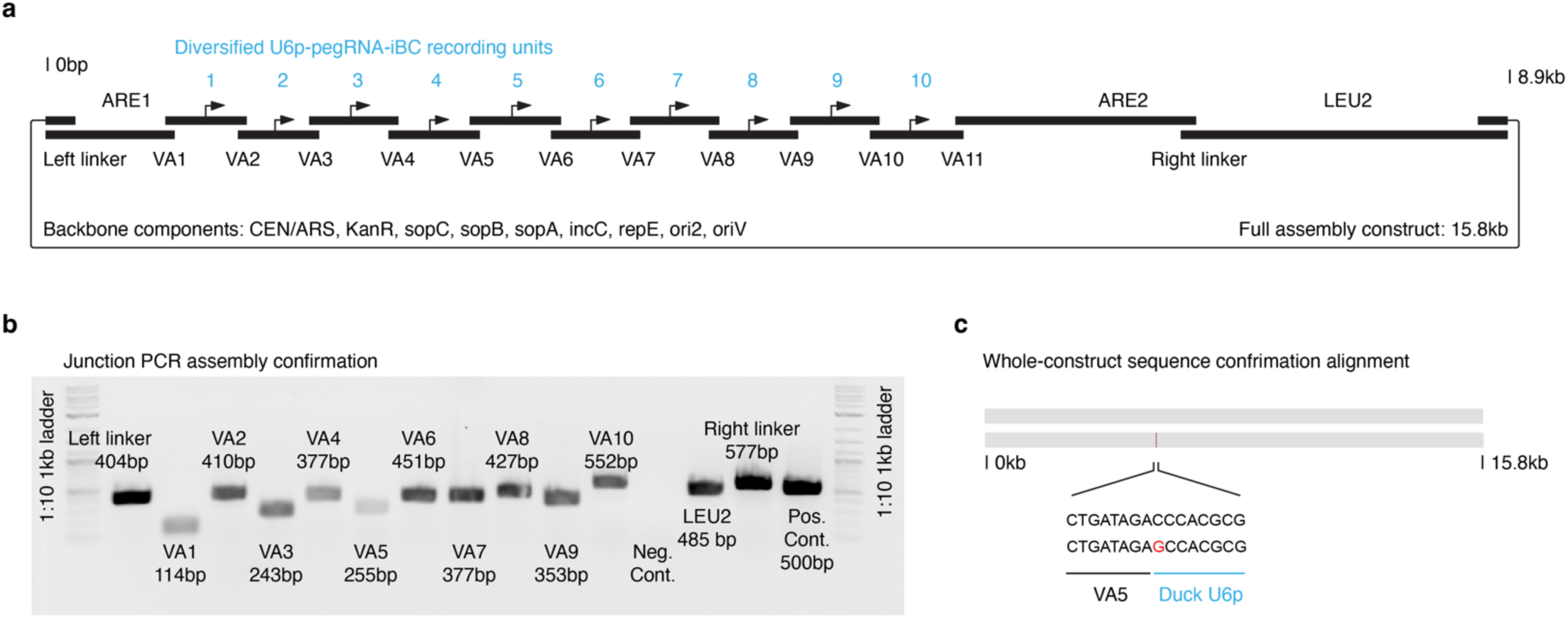
Design and assembly of a 10-unit diversified molecular recording array. **a)** Assembly design showing the 14 fragments flanked with overlapping VEGAS adapters and/or linkers to enable single-step assembly in yeast. VA: vegas adapter, ARE: anti-repressor element, ORI: origin of replication. b) Junction PCR amplicons confirmed effective assembly. **c)** Long-read, whole-plasmid/construct sequencing further confirmed correct assembly. Alignment revealed only a single nucleotide substitution error in the first base pair of the 5th U6 promoter from the Domestic Muscovy Duck *Cairina moschata domestica*. This substitution falls upstream of the four core TFBSs and is not predicted to impact function.

**Figure S13.**
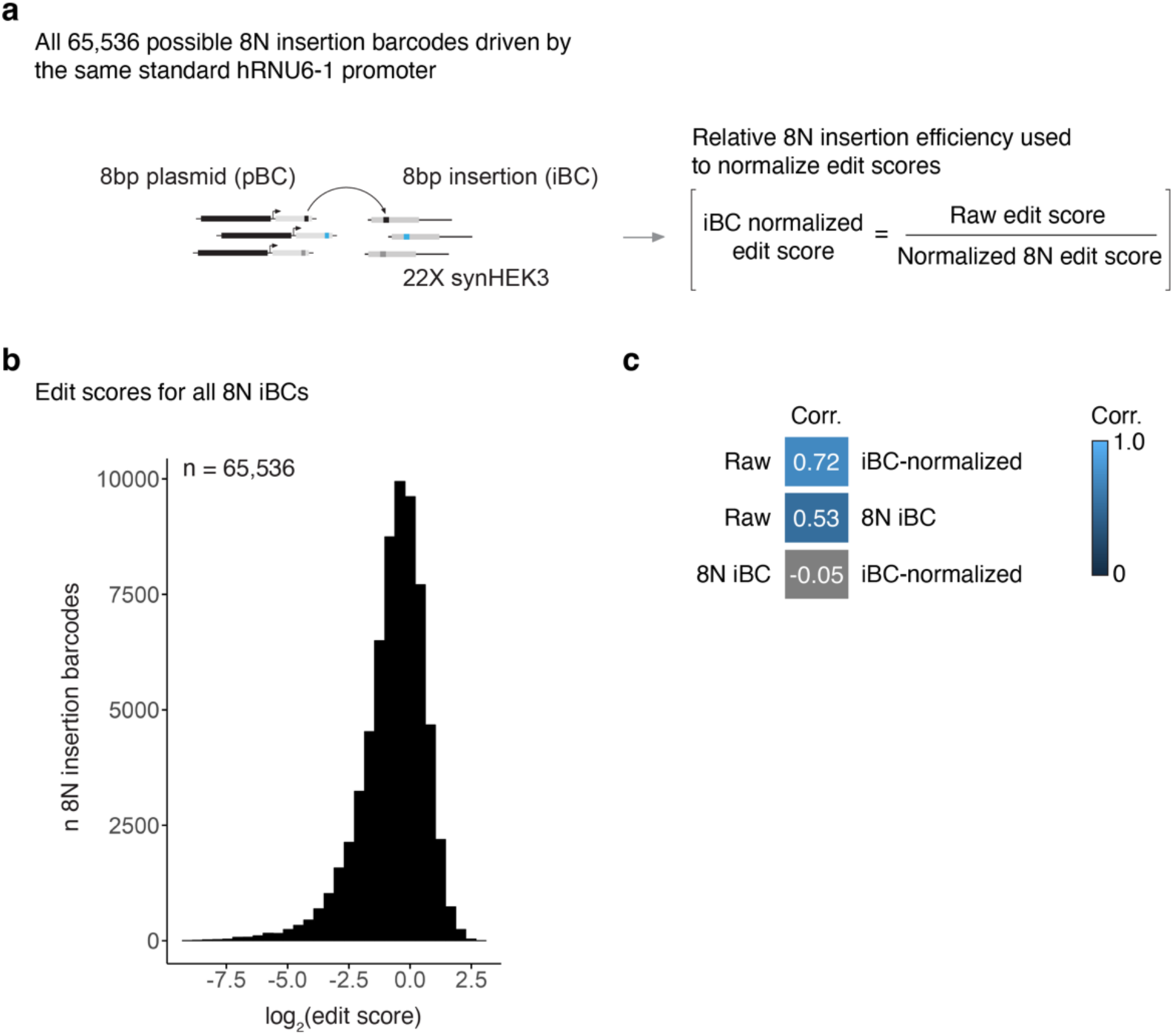
Relative edit scores of all possible 8N insertion barcodes and strategy for edit score normalization. **a)** A library of pegRNAs programmed to insert all 65,536 possible 8N iBCs was driven by the standard human RNU6-1 promoter to assess their relative insertion efficiencies. The resulting iBC edit scores, calculated by dividing each 8 bp sequence’s insertion frequency at *HEK3* by its frequency in the pegRNA library, was used to normalize raw edit scores for diversified U6 promoters paired with a given 8N iBC. **b)** Distribution of edit scores for all 8N iBCs driven by the standard human RNU6-1 promoter. **c)** Correlation between raw edit scores for diversified U6 promoters paired with different 8N iBCs and iBC-normalized edit scores. Accounting for relative 8N iBC insertion efficiency effectively corrected for their influence on diversified U6 promoter edit scores.

**Figure S14.**
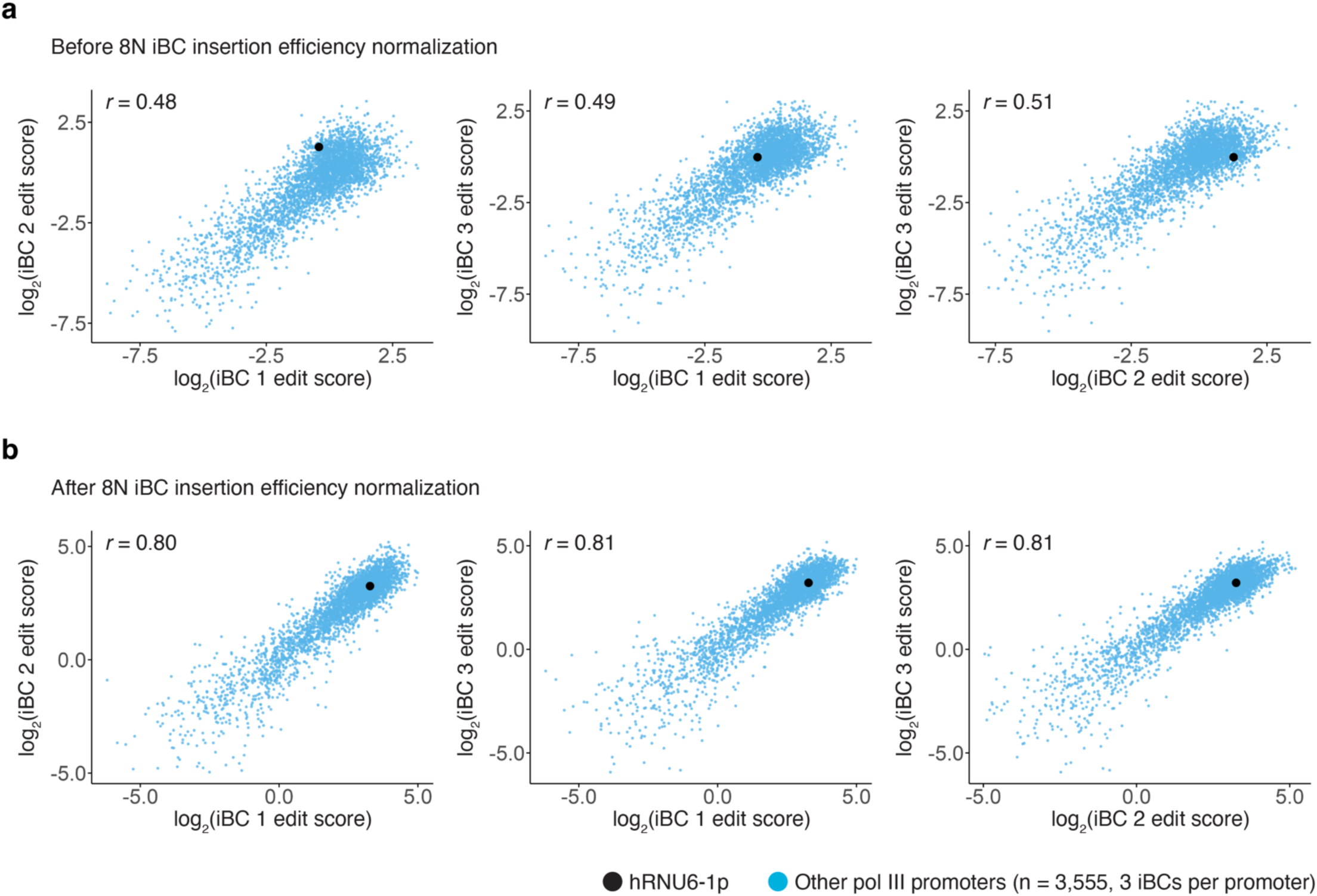
Edit score correlations for independent iBCs across the 3,556 evolutionarily diversified Pol III promoters. **a)** Correlation of raw edit scores across iBCs. **b)** Correlation of edit scores across iBCs following correction for relative barcode insertion efficiencies. Accounting for barcode insertion efficiency substantially improved concordance. In both panels, Pearson correlations, calculated on barcode-normalized edit scores prior to log transformation, are shown.

**Figure S15.**
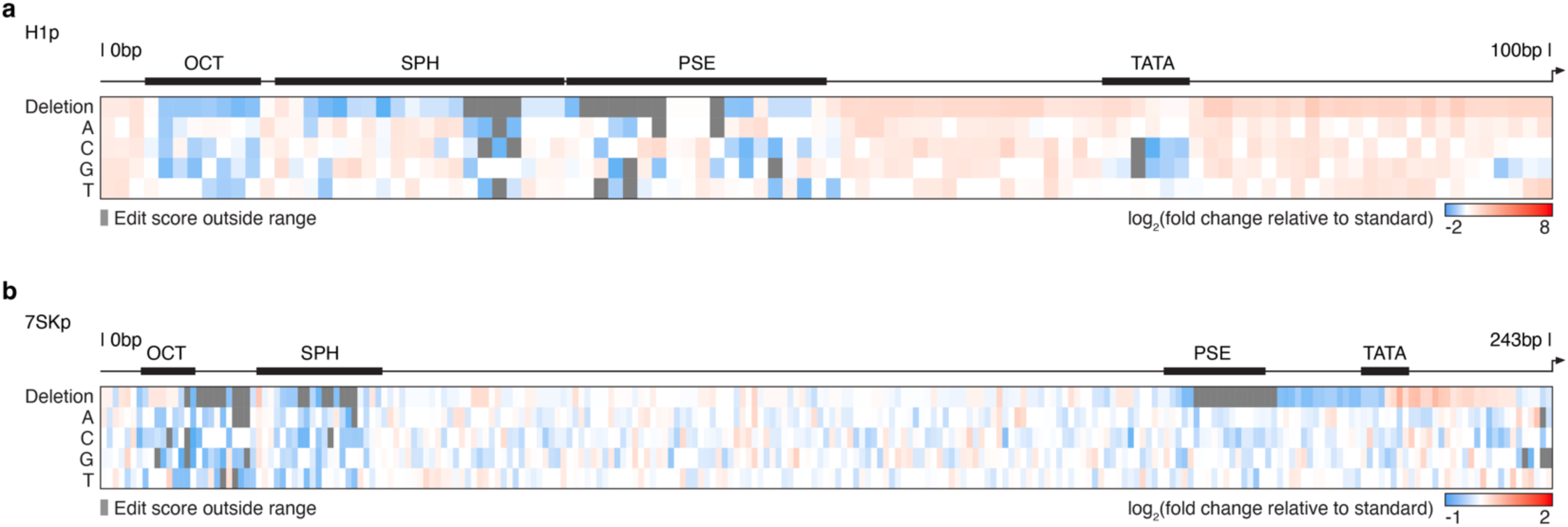
Variant effect maps for human H1 and 7SK promoters. **a)** Variant effect map for the human H1 promoter. **b)** Variant effect map for the human 7SK promoter. Log-transformed fold-changes in edit scores relative to the wildtype human H1 or 7SK promoter are shown. Human H1 and 7SK promoters are relatively tolerant to single nucleotide deletions in the TATA box.

**Figure S16.**
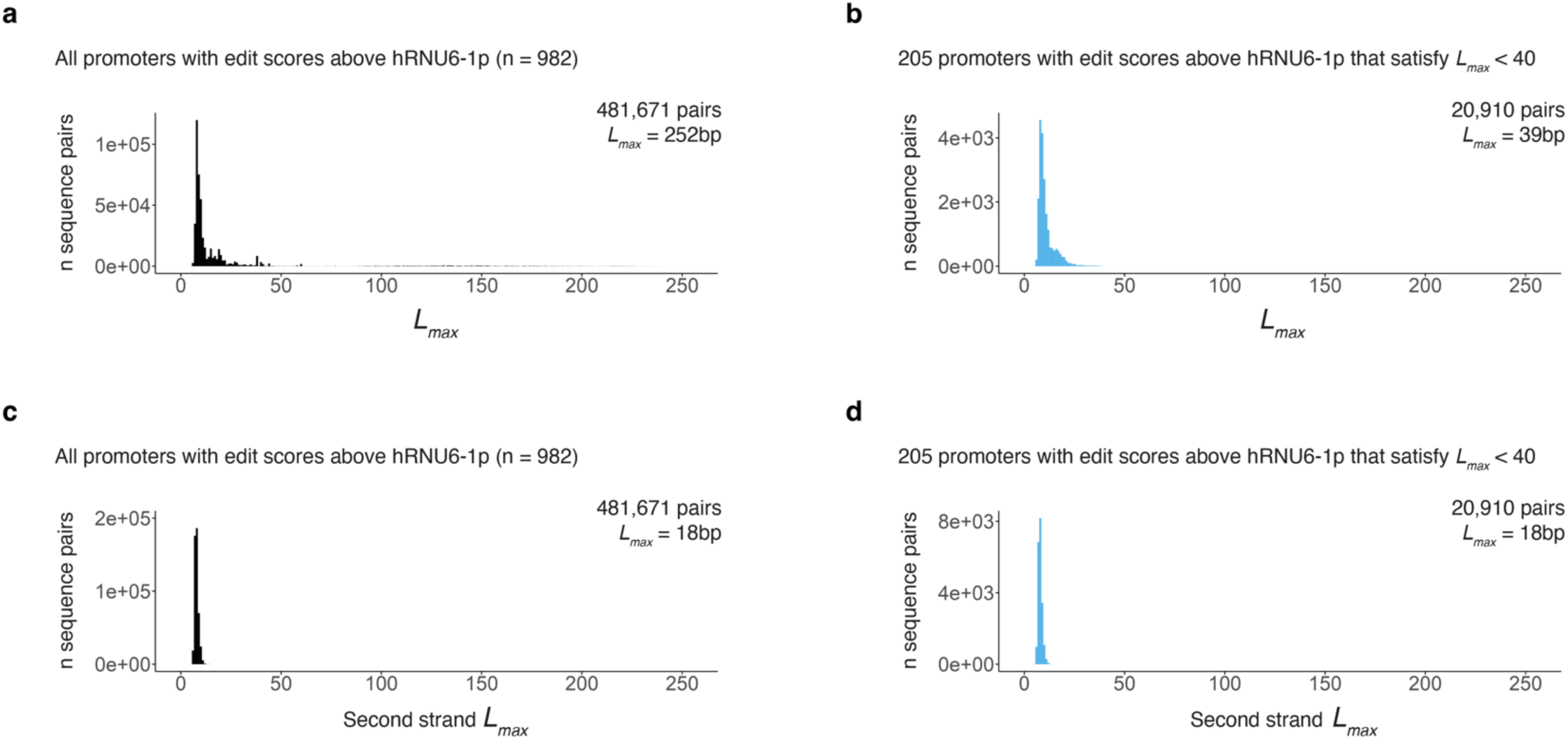
*L_max_* distributions for Pol III promoters with edit scores above standard. **a)** *L_max_* distributions quantifying the maximal shared repeated length between all possible pairs of sequences, in the same orientation, for all Pol III promoters with edit scores above standard (n=982, 481,687 pairs) **b)** *L_max_* distributions depicting how subsets of up to 205 Pol III promoters, in the same orientation, can be used and satisfy *L_max_* < 40, enabling assembly (n=982, 20,910 pairs). **c-d)** *L_max_* distributions as in **a-b**, but for reverse complement comparisons.

**Figure S17.**
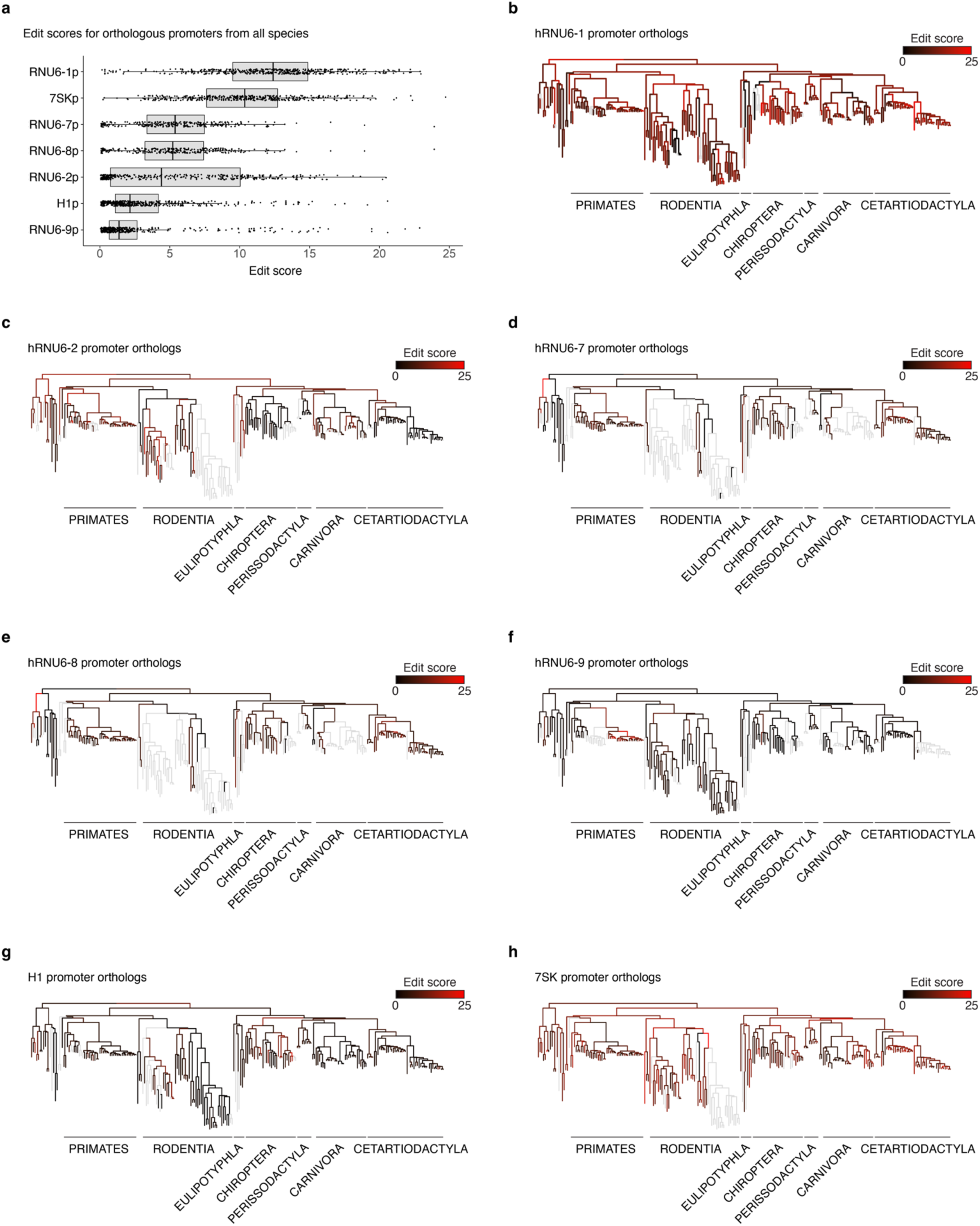
The functional landscape of mammalian Pol III promoter evolution. **a)** Edit scores distributions for ancestral and extant mammalian orthologs of various human Pol III promoters. Boxes represent the 25th, 50th, and 75th percentiles. Whiskers extend from hinge to 1.5 times the interquartile range. **b-h)** Edit scores from panel **a** plotted onto phylogenetic trees to visualize the evolution of functional activity of hRNU6-1p (**b**), hRNU6-2p (**c**), hRNU6-7p (**d**), hRNU6-8p (**e**), hRNU6-9p (**f**), H1p (**g**) and 7SKp (**h**).

## Acknowledgements

We are grateful to members of the Shendure lab and the Seattle Hub for Synthetic Biology for comments, suggestions, and discussions on this work. We are particularly grateful to the Shendure lab gene regulation subgroup for technical advice and deep discussions regarding this work. This work was supported by the Weill Neurohub (to J.S.), the National Institutes of Health (R01HG010632 to J.S.; K99HG012973/R00HG012973 to J.C.; DP5OD036167 to S.P.; RM1HG009491), the Paul G. Allen Frontiers Group (Allen Discovery Center for Cell Lineage Tracing to J.S.), the Brotman Baty Institute for Precision Medicine, and the Seattle Hub for Synthetic Biology, a collaboration between the Allen Institute, the Chan Zuckerberg Initiative (award number CZIF2023-008738), and the University of Washington. T.A.M was supported by a Banting Postdoctoral Fellowship from the Natural Sciences and Engineering Research Council of Canada (NSERC). M.L.T was supported by an award from the Weill Neurohub. H.K. is a Washington Research Foundation Postdoctoral Fellow. J.B.L is a Fellow of the Damon Runyon Cancer Research Foundation (DRG-2435-21). J.S. is an Investigator of the Howard Hughes Medical Institute.

## Author contributions

Conceptualization, T.A.M., and J.S.; Investigation, T.A.M., M.L.T., W.C., F.M.C, J.C, H.L., X.L., H.K., J.B.L., T.L., J.F.N., B.K.M., J.K., A.L.V.C., J.M.G, S.P., and J.S.; Data Curation, T.A.M., and M.L.T.; Formal Analysis, T.A.M., W.C., and M.L.T.; Visualization, T.A.M., and M.L.T.; Resources, J.S.; Supervision, J.S.; Writing – Original Draft, T.A.M., M.L.T., J.S.; Writing – Review & Editing, T.A.M., M.L.T., W.C., F.M.C, J.C, H.L., X.L., H.K., J.B.L., T.L., J.F.N., B.K.M., J.K., A.L.V.C., J.M.G, S.P., and J.S.; Funding Acquisition, J.S.

## Data availability

Raw sequencing data have been uploaded on Sequencing Read Archive (SRA) with associated BioProject ID PRJNA1161643 (https://www.ncbi.nlm.nih.gov/bioproject/PRJNA1161643). Processed data, analysis and visualization code are available at GitHub (https://github.com/shendurelab/Diversified_Parts/), together with construct maps and custom sequencing amplicons used in this work.

## Code availability

Analysis and visualization code are available at GitHub (https://github.com/shendurelab/Diversified_Parts/), together with construct maps and custom sequencing amplicons used in this work.

## Supplementary Tables

**Table S1** - Diversified U6 promoter sequences, paired insertion barcodes, and edit scores.

**Table S2** - Diversified pegRNA scaffold sequences, paired insertion barcodes, and edit scores.

**Table S3** - Saturation mutagenesis U6p-pegRNA cassette sequences, paired insertion barcodes, and edit scores.

**Table S4** - 10-unit diversified recording assembly sequences, predicted and observed activity.

**Table S5** - Zoonomia Pol III promoter sequences, paired insertion barcodes, and edit scores.

**Table S6** - Primer sequences.

## Methods

### Library design and cloning

#### U6 promoter libraries

Promoter sequences of vertebrate orthologs of known transcriptionally active human U6 small nuclear RNA genes were obtained from the ENSEMBL database^74,75^. Sequences were selected for diversity initially using the distance metric from hierarchical clustering (Clustal Omega multiple sequence alignment)^76^ followed by *L_max_* calculations (detailed below) to ensure all promoters satisfied *L_max_* < 40. Additional, sufficiently diverse U6 promoters from vertebrate species (n=4)^24^, transcriptionally active human U6 promoters (n=3)^33^, as well as the canonical human RNU6-1 promoter were also included. Synthetically diversified hRNU6-1 promoters were generated by shuffling nucleotides between the core TFBSs (OCT, SPH, PSE and TATA box) using custom R scripts^77–81^. In a subset of cases, we consulted TFBS profiles from the JASPAR database^82,83^ and further introduced putatively tolerated variants into TFBSs and/or random 3bp spacers sequences between sites (spacers are present in other transcriptional active human U6 promoters) to further increase diversity (n = 52 variants via non-TFBS sequence permutation, n = 30 variants via non-TFBS sequence permutation and SPH TFBS mutation, n = 30 variants via non-TFBS sequence permutation and introduction of random 3bp spacer sequence between the OCT and SPH TFBSs). U6 promoter-pegRNA-pBC cassettes were ordered as eBlocks (IDT) with flanking BsaI golden gate assembly sites^84^. BsaI restriction sites were removed from promoter sequences where required to enable cloning. Also where required, 5bp buffer sequences were inserted flanking the 5’ restriction site to enable commercial synthesis. U6p-pegRNA-pBC eBlocks were then pooled, BsaI digested and ligated (NEB, Cat. No. R3733L) at a 2:1 insert:vector ratio into a minimal backbone^39^ (Twist Bioscience). Cloned libraries were then electroporated into NEB 10-beta electrocompetent E. coli (NEB, Cat. No. C3020K), cultured at 30°C overnight and prepared using a Zymo Pure II (Cat. No. D4200) kit following manufacturer protocols. Singleton validation constructs were confirmed via whole-plasmid sequencing (Primordium Labs). Primer sequences are provided in **Table S6**. U6p-pegRNA-pBC sequences are provided in **Table S1**.

#### pegRNA scaffold libraries

Diversified pegRNA sequences containing complementary R:AR replacement and extension variants (**Fig. 2a**) were generated and paired with respective 5N pBCs using custom R scripts^77–81^. Diversified pegRNA-pBC cassettes were ordered as oligo pools (IDT) with flanking BsaI sites. Oligos were double stranded across multiple low cycle PCR reactions using Q5 polymerase (NEB, Cat. No. M0492L; cycling conditions: 98°C for 30 seconds, 5 cycles of 98°C × 10 seconds, 65°C x 15 seconds and 72°C x 30 seconds). PCR products were then pooled and purified using 2.0x AMPure XP beads (Beckman Coulter, Cat. No. A63880), then BsaI digested and ligated (NEB, Cat. No. R3733L) into a backbone with the standard hRNU6-1 promoter for expression. Plasmid library DNA was prepared as above. Primer sequences are provided in **Table S6**. Diversified pegRNA-pBC sequences are provided in **Table S2**.

#### Miniaturized hRNU6-1p saturation mutagenesis libraries

Saturation mutagenesis variant sequences of the miniaturized hRNU6-1p cassette (**Fig. 3**) were generated and paired with respective 5N pBCs using custom R scripts^77–81^. For the initial deletion series experiments (**Fig. 3a,b**), miniaturized U6p-pegRNA cassettes were ordered as eBlocks with four independent iBCs and cloned as described above for the diversified U6 promoter libraries. Saturation mutagenesis pBC cassettes were ordered as oligo pools (IDT) with flanking BsaI sites. Oligos were double stranded across multiple low cycle PCRs using Q5 polymerase (NEB, Cat. No. M0492L; cycling conditions: 98°C for 30 seconds, 5 cycles of 98°C × 10 seconds, 65°C x 15 seconds and 72°C x 30 seconds). PCR products were then pooled and purified using 2.0x AMPure XP beads (Beckman Coulter, Cat. No. A63880), then BsaI digested and ligated (NEB, Cat. No. R3733L) into a minimal backbone. Plasmid library DNA was prepared as above. Primer sequences are provided in **Table S6**. Diversified min. hRNU6-1p-pegRNA-BC sequences are provided in **Table S3**.

#### Orthologous Pol III promoters (Zoonomia), H1 and 7SK saturation mutagenesis libraries

To select orthologous sequences, we leveraged the cactus alignment (2020v2) from the Zoonomia consortium^57^, relying on the Hal suite of tools^85^.

Briefly, HalLiftover (cactus-bin-v2.7.1) was used with the human interval (hg38) of the Pol 3 promoter as query sequence to all 241 extant mammalian genomes and their reconstructed ancestral sequences (options: --bedType 4 --noDupes 241-mammalian-2020v2.hal). The resulting possibly discontiguous output orthologous intervals were then merged with stitchHalFrags_v2^86^ (modified), requiring that the final interval was within 0.5 to 1.5 fold in length compared to the original query interval size. Sequences not meeting this size threshold were discarded from downstream analysis. In cases for which the intervals spanned different contigs, the sequences were also discarded to avoid complications.

The nucleotide sequences were then obtained from the merged bed file using bedtools (version 2.29.2) getfasta with options -s -fi using the hal genome of the corresponding target species. All resulting sequences with one or more undetermined bases (N) within the queried orthologous region were discarded. Both orientations of remaining orthologous sequences were then pairwise aligned (Biostrings 2.62.0, pairwiseAlignment, options: type=“global”, gapOpening = −2, gapExtension = −8) to their human counterpart to determine correct orientation. The final orientation considered was the one with the largest alignment score to the starting human sequence. After the various filters, out of 481 possible extant and ancestral reconstructed genomes, 437 H1, 426 7SK, 454 U6, 358 RNU6_2, 285 RNU6_7, 286 RN6_8, and 340 RNU6_9 promoter sequences were obtained. Resulting promoters were paired with respective 8N pBCs using custom R scripts^77–81^.

Saturation mutagenesis variant sequences of the human H1 and 7SK promoters were generated and paired with respective 8N pBCs using custom R scripts^77–81^.

Pol III promoter - scaffold - iBC cassettes were ordered as 500bp oligos from Twist with flanking dial out PCR primers^87^ for double stranding and isolation as well as BsaI restriction sites for cloning. BsaI restriction sites were removed from promoter sequences where required to enable cloning. Buffer sequence was added 5’ to the first BsaI site for shorter promoter sequences to assure equivalent size during low-cycle double stranding and subpool isolation PCR (prior to removal following BsaI digestion/cloning). Oligos were double stranded across multiple low cycle PCRs using Q5 polymerase (NEB, Cat. No. M0492L; cycling conditions: 98°C for 30 seconds, 5 cycles of 98°C × 10 seconds, 65°C x 15 seconds and 72°C x 30 seconds). PCR products were then pooled and purified using 2.0x AMPure XP beads (Beckman Coulter, Cat. No. A63880), then BsaI digested and ligated (NEB, Cat. No. R3733L) into a minimal backbone. Plasmid library DNA was prepared as above. Primer sequences are provided in **Table S6**. Pol III promoter and pBC sequences are provided in **Table S5**.

#### 10-unit diversified molecular recording array design and assembly

Top 10 diversified U6 promoter and pegRNA scaffold sequences were paired in reverse rank order (1st promoter with 10th scaffold, 2nd with 9th, etc.). Top parts were selected based on the highest median edit score across cellular contexts. For diversified scaffolds the edit scores from the second barcode pool were used. These Top 10 promoter-scaffold pairings were further assigned specific “<u>GGA</u>NNN” DNA Typewriter iBCs^31^ using custom R scripts (**Fig. 5**). The resulting diversified U6p-pegRNA-iBC units were paired with flanking VEGAS adapters^52^ (10 units, 11 VEGAS adapters) and ordered as sequence-verified double-stranded fragments from STOMICS (**Supplementary Fig. 12**). Additional segments containing auxiliary sequences (ARE, ITRs, etc.) and left/right backbone linkers were also ordered as sequence-verified double-stranded fragments from STOMICS (**Supplementary Fig. 12**). Upon arrival, fragments were amplified using Q5 polymerase, size-verified on a 1% agarose gel, and purified using a Zymo clean and concentrate kit (Cat. No. D4013). Primer sequences are provided in **Table S6**. To generate the backbone fragment, 1 µg of the vector backbone (pSP0769) was linearized using PmeI (NEB, Cat. No. R0560S) for 1h and gel purified.

All resulting fragments were transformed into yeast (*Saccharomyces cerevisiae*) for single-step assembly using the following protocol: 1. The yeast strain BY4741 was grown overnight in 5 mL of 2% YPD media (1% yeast extract, 2% peptone, and 2% dextrose). 2. 1 mL of the overnight yeast culture was transferred to 20 mL of 2% YPD and cultivated for 4 hours at 30°C and 200 rpm. 3. The cells were harvested at 300g for 3 minutes and washed with 20 mL of water. 4. The cells were harvested again and washed with 0.1M Lithium Acetate (LiAc). 5. The cells were harvested, and the supernatant was removed. The cell pellet was then resuspended in 0.1M LiAc that remained in the tube. 6. The cells were transferred to a 1.5 mL tube, harvested at 300g for 3 minutes, resuspended in 200 µL of 0.1M LiAc, and kept on ice. 7. The segments and linearized vector were pooled together at a concentration of approximately 0.5 pmol each. 8. The transformation mix was prepared by combining 240 µL of 44% polyethylene glycol (PEG) solution, 36 µL of 1M LiAc, and 25 µL of herring sperm DNA. 9. 20 µL of cells were transferred to the tube containing the pooled DNA and vortexed briefly. 10. The transformation mix was added to the DNA + cells solution, and the mixture was vortexed at high speed for 10 seconds. 11. The mixture was transferred to a 30°C incubator with rotation and left for 30 minutes. 12. 36 µL of DMSO was added to the tube, followed by a 15-minute incubation at 42°C using a water bath. 13. Cells were harvested and resuspended in 200 µL of 5 mM CaCl₂ before being plated onto SC -LEU plates. 14. Plates were incubated at 30°C, and the presence of colonies was checked after 2-3 days.

Candidates were initially checked through junction PCR, where the presence of each junction between all the transformed segments was verified using segment-specific primer pairs (**Supplementary Fig. 12**). Primer sequences are provided in **Table S6**. Yeast cells that passed this initial check were grown in 5 mL of SC -LEU media overnight, and the plasmids were extracted using the yeast miniprep I kit from Zymo Research (Cat. No. D2001). The plasmids were then transformed into E. coli (EPI300 cells) through electroporation. E. coli cells were subjected to a miniprep (Zymo kit), and the 10-unit assembly construct was sequence-verified through commercial long-read sequencing (Plasmidsaurus-nanopore). The final construct presented only a single SNP at the first base of U6 promoter number 5 (**Supplementary Fig. 12**).

#### *L_max_* calculations

To calculate Lmax, we wrote a pipeline that takes as input a list of sequences and first generates a dataframe containing all possible pairs of sequences in the forward and reverse orientation: n possible pairs = (n x n-1)/2 in each orientation. The pipeline iterates through each row of the sequence-pair dataframe applying a longest common substring function^88^ to return the length and identity of the longest shared sequence repeat in each pair of sequences in a given set (**Supplementary Fig. 1**). The resulting *L_max_* distributions can be filtered to select sets of sequences that satisfy any *L_max_* threshold (e.g. *L_max_* < 40 used here) (**Supplementary Fig. 1**).

### Cell Lines and Culture

#### K562 cell culture

K562 cells (ATCC Cat. No. CCL-243)^89^ were grown with 5% CO_2_ at 37°C and cultured in RPMI 1640 + L-Glutamine (GIBCO, Cat. No. 11-875-093) supplemented with 10% fetal bovine serum (Rocky Mountain Biologicals, Cat No. FBS-BSC) and 1% penicillin-streptomycin (Thermo Fisher Scientific, Cat. No. 15070063).

#### HEK293T cell culture

HEK293T cells (ATCC Cat. No. CRL-11268) were grown with 5% CO_2_ at 37°C and cultured in high glucose DMEM (GIBCO, Cat. No. 11965092) supplemented with 10% fetal bovine serum (Rocky Mountain Biologicals, Cat No. FBS-BSC) and 1% penicillin-streptomycin (Thermo Fisher Scientific, Cat. No. 15070063).

#### iPSC culture

WTC11 iPSCs^90^ were grown with 5% CO_2_ at 37°C cultured in mTeSR Plus Basal Medium (Stemcell technologies; Cat. No. 100-0276) on Greiner Cellstar plates (Sigma-Aldrich; assorted Cat. Nos.) coated with Geltrex™ LDEV-Free, hESC-Qualified, Reduced Growth Factor Basement Membrane Matrix (Gibco; Cat. No. A1413302) diluted 1:100 in Knockout DMEM (GIBCO/Thermo Fisher Scientific; Cat. No. 10829018). Cells were passaged by washing cells with PBS (GIBCO/Thermo Fisher Scientific; Cat. No. 10010023), dissociating with StemPro Accutase Cell Dissociation Reagent (GIBCO/Thermo Fisher Scientific; Cat. No. A1110501) and resuspending cell pellets in mTeSR Plus Basal Medium supplemented with 0.1% dihydrochloride ROCK Inhibitor (Stemcell technologies; Cat. No. Y-27632). mTeSR plus media was replaced every other day.

#### mESC culture

E14 mESCs were grown with 5% CO2 at 37°C cultured in media composed of Advanced DMEM (Gibco, cat. no. 11965118) supplemented with 15% KSR (Gibco, cat. no. 10828028), 1X NEAA (Gibco, cat. no. 11140050), 1X Glutamax (Gibco, cat. no. 35050061), 1 mM sodium pyruvate (Gibco), 0.5 µM 2-Mercaptoethanol (ThermoFisher, cat. no. 31350010), and 1000 U/ml LIF (ESGRO) on 6cm dishes that had been pre-coated with 0.2% gelatin (Millipore Sigma, cat. no. G1890). For passaging, cells were dissociated with 0.05% Trypsin-EDTA (Gibco, cat. no. 25300120), pipetted gently to generate a single-cell suspension, then the trypsinization reaction was quenched with a wash medium composed of Advanced DMEM/F-12 (Gibco, cat. no. 12634010) supplemented with 5% FBS (Cytiva, cat. no. SH30071.03HI) before resuspending in culture media. Culture media was replaced every day.

### Cell line generation

#### K562

The monoclonal PE2-K562 cell line was generated using piggyBac transposition. Specifically, 500ng of a PE2 cargo construct^91^ and 100ng of a super piggyBac transposase expression vector (System Biosciences, Cat. No. PB210PA-1) were mixed and transfected using lipofectamine 3000 (Thermo Fisher Scientific; Cat. No. L3000015) following manufacturer protocol. PE2 expressing cells were then selected by antibiotic resistance (puromycin), single cell-sorted into 96-well plates using a flow sorter and cultured for 2-3 weeks until confluency. Multiple lines were then tested for prime editing insertion efficiency using a pegRNA expression construct programmed to insert “CTT” at the *HEK3* locus (Addgene #132778)^38^ and the line with the highest editing efficiency was selected for use. The 22x*synHEK3* K562 line was generated using piggyBac transposition and mapped with a T7-promoter strategy as previously described^49^.

#### HEK293T

The polyclonal PEmax-HEK293T cell line was generated using piggyBac transposition. Specifically, a PEmax cargo construct and super piggyBac transposase expression vector (System Biosciences, Cat. No. PB210PA-1) were mixed and transfected using lipofectamine 3000 (Thermo Fisher Scientific; Cat. No. L3000015) at a 5:1 molar ratio following manufacturer protocol. PEmax expressing cells were then selected by antibiotic resistance (blasticidin) and PEmax expression was confirmed by fluorescence. The polyclonal 6xTAPE-PEmax-HEK293T line was generated using piggyBac transposition into the previously generated HEK293T-PEmax line. Specifically, 2,160ng of 6xTAPE construct, and 240 ng of a super piggyBac transposase expression vector (System Biosciences, Cat. No. PB210PA-1) were mixed and transfected using lipofectamine 3000 (Thermo Fisher Scientific; Cat. No. L3000015) following manufacturer protocol and then selected with 400 ug/mL of hygromycin for 1 week.

#### iPSC

The monoclonal PEmax WTC11 iPSC line was generated by piggyBac transposition. Specifically, a PEmax cargo construct and a super piggyBac transposase expression vector (System Biosciences, Cat. No. PB210PA-1) were mixed at a 5:1 molar ratio and nucleofected using the CB-150 program and P3 primary reagents (Lonza, Cat. No. V4XP-3032) on a Lonza 4D nucleofector following manufacturer protocol. PEmax expressing cells were then selected by antibiotic resistance (blasticidin), single cell-sorted using limiting dilution, and cultured for 2-3 weeks until confluent. Multiple lines were tested for 5N prime editing insertion efficiency at the *HEK3* locus and the line with the highest editing efficiency was selected for use.

#### mESC

The monoclonal PEmax-*synHEK3* E14 mESC line was generated by piggyBac transposition. Specifically, a PEmax cargo construct, a *synHEK3* cargo construct, and a super piggyBac transposase expression vector (System Biosciences, Cat. No. PB210PA-1) were mixed at a 17:2:1 molar ratio (85%, 10%, 5%) and transfected using lipofectamine 2000 (Thermo Fisher Scientific; Cat. No. 11668027) following manufacturer protocol. PEmax-expressing cells were then selected by antibiotic resistance (puromycin) for seven days, then the top 10% of GFP+ cells were sorted into a single-cell suspension. These sorted cells were plated on a feeder layer of mitotically inactive mouse embryo fibroblasts (MEFs) to grow into colonies. Monoclonal colonies were then handpicked and further expanded and frozen for future use. The number of integrated *synHEK3* target sites was estimated using diverse barcodes paired with each *synHEK3* target construct that were prepared sequenced as part of the target amplicon library (see below).

### Transfection

#### K562

All libraries were transfected using lipofectamine 3000 (Thermo Fisher Scientific; Cat. No. L3000015) following manufacturer’s specifications. 1.5 × 10^5^ cells were seeded the day prior to transfection. 500 ng of each library was then mixed with 100 ng of a GFP co-transformation marker (pmaxGFP, Lonza) and transfected in triplicate or quadruplicate in 24 well plates. Genomic DNA was harvested from cells 3-4 days after transfection. For the 8N iBC and Zoonomia Pol III promoter libraries, 1 × 10^6^ cells were nucleofected with 1000 ng library, 1000 ng of a PEmax construct, and 250 ng of a GFP co-transformation marker (pmaxGFP, Lonza) using a Lonza 4D nucleofector (Lonza; Cat. No. V4SC-2096) in quadruplicate following manufacturer’s specifications.

#### HEK293T

All libraries were transfected using lipofectamine 3000 (Thermo Fisher Scientific; Cat. No. L3000015) following manufacturer’s specifications. 3 × 10^5^ cells were seeded the day prior to transfection. For the U6 promoter library, 1000 ng of each library was then mixed with 250 ng of a GFP co-transformation marker (pmaxGFP, Lonza) and transfected across 8 wells of a 12 well plate. For the diversified pegRNA and saturation mutagenesis libraries, 500 ng of each library was then mixed with 100 ng of a GFP co-transformation marker (pmaxGFP, Lonza) and transfected in triplicate or quadruplicate in 24 well plates. Genomic DNA was harvested from cells 3 days after transfection. The 10-unit assembly was delivered via transfection with lipofectamine 3000 (Thermo Fisher Scientific; Cat. No. L3000015) following manufacturer’s specifications. 3 × 10^5^ cells were seeded the day prior to transfection. 500 ng of the 10-unit assembly was then mixed with 300 ng of a PEmax construct, 125 ng of a GFP co-transformation marker (pmaxGFP, Lonza), and 100 ng of a super piggyBac transposase expression vector (System Biosciences, Cat. No. PB210PA-1) and transfected across 4 wells of a 24 well plate.

#### iPSCs

All libraries were nucleofected using a Lonza 4D nucleofector following manufacturer’s specifications. iPSCs were dissociated and resuspended in mTeSR plus basal media supplemented with ROCKi. 2.2 × 10^5^ cells were nucleofected with 2000 ng of each library, 1000 ng of PEmax construct, and 500 ng of pmaxGFP co-transformation marker (Lonza) using P3 reagents and the CB-150 program on the Lonza 4D nucleofector. Four replicate nucleofections per library were then plated into separate Geltrex coated wells of a 24 well plate. Genomic DNA was harvested from cells 3-4 days after nucleofection.

#### mESCs

All libraries were transfected using lipofectamine 2000 (Thermo Fisher Scientific; Cat. No. 11668019) following manufacturer’s specifications. Transfection reagents were mixed with DNA (1300 ng of library and 145 ng of PEmax per replicate) and allowed to incubate for 20 minutes. During this time, 3.6 × 10^5^ freshly dissociated cells were plated into each of four gelatin-coated wells of a 12 well plate. Transfection reagents were then added to the cells while still in suspension. The plate was then placed in the incubator at 37°C at 5% CO2 and gently rocked across its horizontal and vertical axis to evenly plate the cells. Media was changed the day after transfection. Genomic DNA was harvested from cells 3 days after transfection.

### Genomic DNA extraction

Genomic DNA was extracted as follows: Harvested cells were washed with PBS, then 200 µl of freshly prepared lysis buffer (10 mM Tris-HCl, pH 7.5; 0.05% SDS; 25 µg/ml protease (Thermo Fisher Scientific, Cat. No. EO0491)) per 0.5-1M cells was added directly into each well of the tissue culture plate. The genomic DNA mixture was then incubated at 50°C for 1 h, followed by a 30 min 80°C enzyme inactivation step.

### Library preparation and sequencing

#### Plasmid barcode amplicon sequencing library preparation

pBC amplicon sequencing libraries were generated using a two step PCR process to amplify barcodes then append sequencing adapters and sample indices. pBCs were amplified using a forward primer that binds the gRNA scaffold (U6 promoters), the hRNU6-1 promoter (pegRNA libraries), or upstream of the miniaturized U6 promoter (saturation mutagenesis miniaturized hRNU6-1p-pegRNA libraries) along with a universal reverse primer that binds the plasmid backbone. Plasmid libraries were amplified using Q5 polymerase in quadruplicate (NEB, Cat. No. M0492L; cycling conditions: 98°C for 30 seconds, 15 cycles of 98°C × 10 seconds, 65°C x 15 seconds and 72°C x 40 seconds). SYBR Green (Thermo Fisher Scientific, Cat. No. S7567) was added to track the amplification curve. PCR products were pooled and purified using 1.2x AMPure XP beads (Beckman Coulter, Cat. No. A63880). Sequence flow cell adapters and dual sample indices were then appended in the second PCR reaction using Q5 polymerase (NEB, Cat. No. M0492L; cycling conditions: 98°C for 30 seconds, 5 cycles of 98°C × 10 seconds, 65°C x 15 seconds and 72°C x 30 seconds). PCR products were purified using 0.9x AMPure XP beads (Beckman Coulter, Cat. No. A63880) and assessed on an Agilent 4200 TapeStation before sequencing. Primer sequences are provided in **Table S6**.

#### *HEK3* locus and *synHEK3* amplicon sequencing library preparation

The *HEK3* and *synHEK3* target loci amplicon sequencing libraries were generated using a similar two step PCR process to amplify targets then append sequencing adapters and sample indices. 2ul of cell lysate was used as input to a 50ul PCR reaction using KAPA Robust polymerase (KAPA Biosystems, Cat. No. 2GRHSRMKB; cycling conditions: 95°C for 3 min, 22-29 cycles of 95°C x 15 seconds, 65°C x 15 seconds and 72°C x 30 seconds). SYBR Green (Thermo Fisher Scientific, Cat. No. S7567) was added to track the amplification curve. PCR products were pooled and purified using 1.2x AMPure XP beads (Beckman Coulter, Cat. No. A63880). Sequence flow cell adapters and dual sample indices were then appended in a second PCR reaction (cycling conditions: 98°C for 30 seconds, 5 cycles of 98°C × 10 seconds, 65°C x 15 seconds and 72°C x 30 seconds). PCR products were purified using 0.9x AMPure XP beads (Beckman Coulter, Cat. No. A63880) and assessed on an Agilent 4200 TapeStation before sequencing. Primer sequences are provided in **Table S6**.

Libraries were sequenced on an Illumina MiSeq sequencer, lllumina NextSeq500 sequencer, or Illumina NextSeq2000 sequencer following manufacturer’s protocol.

#### Edit score calculations and insertion barcode normalization

pBC and iBC counts were extracted from plasmid library and *HEK3* locus sequencing reads using pattern matching functions. Specifically, we required a perfect match to the 15bp spanning the intended 5N barcode and sequences flanking the edit site in the RTT and PBS in plasmid and edited read datasets to count a barcode. For 8N barcodes this was extended to 18bp. pBC and iBC frequencies were then calculated for each library, and the raw edit score was calculated as iBC freq. / pBC freq. for each replicate. Raw edit scores were divided by the normalized insertion efficiency of the paired barcode to correct for insertion barcode efficiency (**Supplementary Figs. S2**, **S13**). Correlations between cellular contexts were calculated on the barcode-normalized edit scores. Note we initially selected and tested an additional four promoters and two scaffolds with hierarchical clustering as the diversity metric that ultimately did not satisfy the more stringent criterion of *L_max_* < 40 and so were removed from analysis and final reported functional part sets. Any part with an edit score below 0.005 was assigned an edit score of zero in final results tables (**Tables S1-S3**).

